# A complementary learning system for continual episodic memory in large language models

**DOI:** 10.64898/2026.08.24.746712

**Authors:** Xu Pan, Ely Hahami, Roy Siegelmann, Haim Sompolinsky

**Author notes:** **For correspondence:** (HS). These authors contributed equally to this work (Xu Pan, Ely Hahami).

## Abstract

Humans retain memories of individual experiences for a lifetime, an ability attributed to a complementary learning system in which a fast process encodes episodes and a slow process integrates them into semantic knowledge. In classical Hebbian models such as Hopfield networks, memory traces are superposed in shared weights. This makes learning naturally continual but causes strong interference among correlated memories, a failure that reappears as catastrophic forgetting in deep networks. Here we use a large language model as a model system for continual episodic memory, with its pretrained weights supplying the semantic context in which new episodes are embedded. Fast learning is implemented by a hippocampus-like module that assigns each episode to a dedicated, extremely sparse low-rank adapter; competitive gating then selects among these separated traces during recall. Across streams of up to 1,000 factual and autobiographical episodes, each adapter requires only 2–3 parameters per token while preserving excellent recall. An internal retrieval-augmented generation mechanism reconstructs the selected episode in context and supports high-accuracy question answering over stored memories. Finally, slow cortical consolidation is modeled by fine-tuning the base weights through batch replay, enabling reconstruction and direct question answering without episodic adapters. Together, fast storage and slow consolidation implement both components of a complementary learning system within a single language model, yielding a neural-network model that stores, recalls, and consolidates naturalistic episodic memories, thereby capturing key functional features of human memory.

## Introduction

Humans continually form memories of individual experiences and retain them over days, years, or a lifetime. Cognitive science attributes this ability to a complementary learning system (CLS) (***McClelland et al., 1995***; ***Kumaran et al., 2016***): a fast process, centered on the hippocampus, that rapidly encodes each new episode, and a slow process in the neocortex that gradually integrates information from episodes into structured semantic knowledge (***Squire, 2004***; ***Tse et al., 2007***).

There is no widely accepted network model of the fast episodic system. The classical candidate is the Hopfield content-addressable memory (CAM) (***Hopfield, 1982***). As a fast memory, CAM has two appealing properties. First, learning is continual by the nature of the Hebbian rule: each new pattern is stored by a local, additive weight update, without revisiting previously stored patterns. Second, its capacity is substantial, and below a critical load stored patterns can be recovered as faithful fixed-point attractors (***Amit et al., 1985***).

The same mechanism also creates its central weakness. Because every memory is stored by overlaying its trace on a shared set of weights, the traces interfere. For random patterns, this interference remains tolerable below the critical load, beyond which retrieval collapses (***Amit et al., 1985***). For correlated patterns, the problem is more severe: the Hebbian rule is highly sensitive to correlations, and capacity collapses well before the random-pattern limit (***Löwe, 1998***). The corresponding failure in modern networks trained sequentially is called catastrophic forgetting (***McCloskey and Cohen, 1989***; ***Ratcliff, 1990***). In both cases, new traces are written over old ones in shared weights. Interference among overlapping traces is therefore a common obstacle to continual episodic memory in associative memories and deep networks.

The brain is thought to reduce this interference through pattern separation. The hippocampus re-represents correlated experiences in a decorrelated form, minimizing overlap among their memory traces (***O’Reilly and McClelland, 1994***; ***Yassa and Stark, 2011***). Sparsity is one proposed mechanism, because sparse representations share fewer active units and therefore interfere less. Effective separation also depends on competition mediated by inhibition. During encoding, inhibitory competition restricts the neurons allocated to a new trace, keeping the resulting engram sparse (***Stefanelli et al., 2016***). During recall, competition allows a cue to activate the matching trace while suppressing competing traces (***Rashid et al., 2016***). Sparse allocation and competitive selection are central to models of memory engrams in the hippocampus and amygdala (***Rao-Ruiz et al., 2019***; ***Kim et al., 2013***; ***Han et al., 2007***). These observations motivate two functional requirements for a continual episodic-memory system: strongly separated traces during storage and competitive selection among traces during recall.

Real human episodic memories, however, are not the binary random patterns of common attractor models of associative memory. They are narratives rich in semantic structure. They relate to existing knowledge, interact with other memories, and may eventually contribute to semantic knowledge. They are also acquired sequentially, creating potential interference both among successive memories and with previously acquired skills. Modeling such memories requires a substrate that already contains broad semantic knowledge, within which a new episode can be embedded and to which it can be related. A pretrained large language model provides such a substrate.

An LLM is not itself a solution to continual episodic memory. Naively fine-tuning it on a stream of episodes overwrites earlier memories and can erode its general capabilities (***Luo et al., 2025***; ***Biderman et al., 2024***). Standard continual-learning remedies do not transfer cleanly to the present setting. Replay and regularization have generally been developed for task-incremental learning rather than episodic streams in which each episode arrives once and earlier episodes are not replayed during subsequent encoding (***Parisi et al., 2019***; ***Kirkpatrick et al., 2017***), while retrieval-augmented generation stores episodes in an external text repository rather than in the network weights (***Lewis et al., 2020***; ***Ovadia et al., 2024***).

Here we use an LLM as a model system for fast episodic memory. Each episode is encoded into a dedicated masked low-rank adapter, so that different episodes occupy nonoverlapping parameter traces. Extreme sparsity controls the cost of each trace, allowing many such memories to coexist within a small parameter budget. Each adapter is stored together with a semantic embedding that serves as its retrieval key. At recall, a cue is embedded and competitive winner-take-all routing selects the adapter with the most similar key. Thus, the present model implements separated traces during storage and competitive selection during recall. It does not model the biological competition that recruits neurons into an engram during its formation.

The framework also includes a slower consolidation process. Stored episodes are replayed in batches to fine-tune the base model, allowing their contents to be incorporated into shared model weights and subsequently accessed without loading an episodic adapter. This provides a functional analog of the transition from fast, episode-specific storage to slower integration with semantic knowledge (***Wilson and McNaughton, 1994***; ***Diekelmann and Born, 2010***).

We evaluate the framework on sequential streams of up to 1,000 factual and autobiographical episodes. The experiments test whether the system can preserve individual episodes without interference, retrieve the appropriate trace from a semantic cue, use recalled episodes for question answering, and consolidate them into the base model through replay. Together, these experiments examine both the capacity of the proposed fast memory and the more difficult transition from faithful episodic recall to knowledge integrated into the model’s shared weights.

A preliminary version of this work was posted on arXiv in April 2025 (Pan et al., 2025b).

## Results

Throughout, we use *episode* for a single experience, such as reading a news article, meeting a colleague, or living through an event, and *passage* for the text that conveys it. Encoding an episode yields an *episodic memory*: a stored trace that a later cue can recall, whether it’s a fact about the world or a personal event. In the classical taxonomy, the former is closer to *semantic* memory, and the latter to *episodic* memory. We focus on the *continual learning* aspect of episodic memory: each episode arrives once in the stream and is encoded before the stream advances, without replay of earlier episodes during subsequent encoding. An episode may nevertheless require multiple optimization steps to encode (Methods). We use *T* for the length of a sequential stream under continual fine-tuning and *N* for the number of stored episodes available to the gated-memory or consolidation system.

### Failures of continual fine-tuning

We ask whether a large language model can sequentially acquire everyday human memories (e.g., news pieces, personal experiences), and retain them faithfully as new episodes continue to arrive. This is a continual learning problem, but one that differs from the typical LLM post-training setting in which a model is adapted to a new skill, such as a discrimination task or response style, in which case the fine-tuning typically involves multiple examples of the new task. Here, each episode arrives once and must be encoded before the stream advances, without replaying earlier episodes. Specifically, we used two datasets representing two types of declarative memory (***Squire, 2004***; ***Tulving, 2002***): a dataset of Wikipedia paragraphs for factual knowledge about the world (the *Wiki* dataset) and a dataset of fictional-character narratives for autobiographical experiences (the *Character* dataset). The Wiki dataset contains 1,000 Wikipedia paragraphs published after the model’s training data cutoff (length 76.8 ± 9.6 tokens, details in Methods). The paragraphs are sampled from a large, diverse pool of Wikipedia entries, ensuring that episodes are largely independent of one another. The Character dataset contains 1,000 first-person episodes (length 95.1 ± 12.7 tokens) spanning 50 years of a single fictional character’s life. The dataset is synthesized by an LLM-based pipeline that maintains cross-episode consistency through a cumulative summary of prior years (details in Methods). Unlike the Wikipedia articles, these episodes are inherently correlated: the same family members, colleagues, and locations recur across decades. In what follows, each *episode* is a single Wiki paragraph or character narrative.

Every experiment below is run with two base LLM models, Llama-3.1-8B-Instruct (***Grattafiori et al., 2024***) and Qwen3-8B (***Yang et al., 2025***). We show Llama in the main figures and Qwen3-8B in Supplementary Note 7 (Supplementary Figs. 5–12); each figure caption names its Qwen counterpart. Every result is qualitatively the same in both models, and we note in the text the few places where they differ.

A defining feature of declarative memory is the verbal recall of episodes given a cue (***Tulving and Thomson, 1973***; ***Cohen and Squire, 1980***; ***Squire and Zola-Morgan, 1991***; ***Tulving, 2002***). We adopt this reconstruction capability as a key performance measure for our experiments and evaluation. We first tested sequential fine-tuning on Llama, training on a sequence of up to 1,000 episodes and measuring how well the model can still recall previously learned episodes after all training is complete. As in other memory systems, we need to specify the nature of the recalling cue. Here we adopt a variant of the pattern completion paradigm where, during recall, the *theme* of one of the episodes is presented and we test whether this triggers the recall of the full episode. Concomitantly, during memorization of the incoming episode, the model is trained to generate its passage conditioned on a prompt containing its *theme*. The themes in the Wiki dataset are the titles of the Wiki articles; the themes in the Character dataset are thematic summarizations of the episodes synthesized along with the passages. Our main evaluation metric is an LLM judge whose scoring prompt is designed to measure the faithfulness of semantic content rather than surface-level lexical overlap. Supplementary Note 6 compares the judge with Levenshtein ratio, ROUGE, and teacher-forced token accuracy, and shows that the qualitative conclusions are unchanged while also illustrating how surface-based and teacher-forced metrics can misrepresent free-generation recall.

The result exhibits severe catastrophic forgetting across both datasets (Fig. 1A,B,D,E): recall of older episodes degrades substantially following the subsequent encoding of only a dozen episodes. Parameter-efficient LoRA has been reported to mitigate catastrophic forgetting (***Hu et al., 2022***; ***Biderman et al., 2024***), but in our setting its benefit is marginal and inconsistent. The only clear gain is all-layer LoRA (0.176% of parameters) on Wiki at *T* = 100, which slows forgetting relative to full fine-tuning (Fig. 1A). On the Character dataset it offers no benefit (Fig. 1D), and restricting the update further to 4 layers (0.022% of parameters) makes forgetting worse. The same qualitative pattern holds at *T* = 10 (Supplementary Fig. 1). Sequential fine-tuning potentially also degrades the model’s general capabilities (Fig. 1C,F). We test this effect by tracking three standard benchmarks, WinoGrande (***Sakaguchi et al., 2021***), HellaSwag (***Zellers et al., 2019***), and MMLU (***Hendrycks et al., 2021***), normalized by the base model’s score before fine-tuning. When using the Llama model, the forgetting of the general capabilities depends on the fine-tuning methods. Full-parameter training of 1,000 memories retains only around 75% of the original score. On the other hand, LoRA-based training is more robust, with both variants remaining above 96% throughout. However, the Qwen3-8B shows minimal to no degradation of general capabilities regardless of the fine-tuning methods (Supplementary Fig. 5), yet its recall forgetting curves are qualitatively the same as the Llama model.

**Figure 1.**
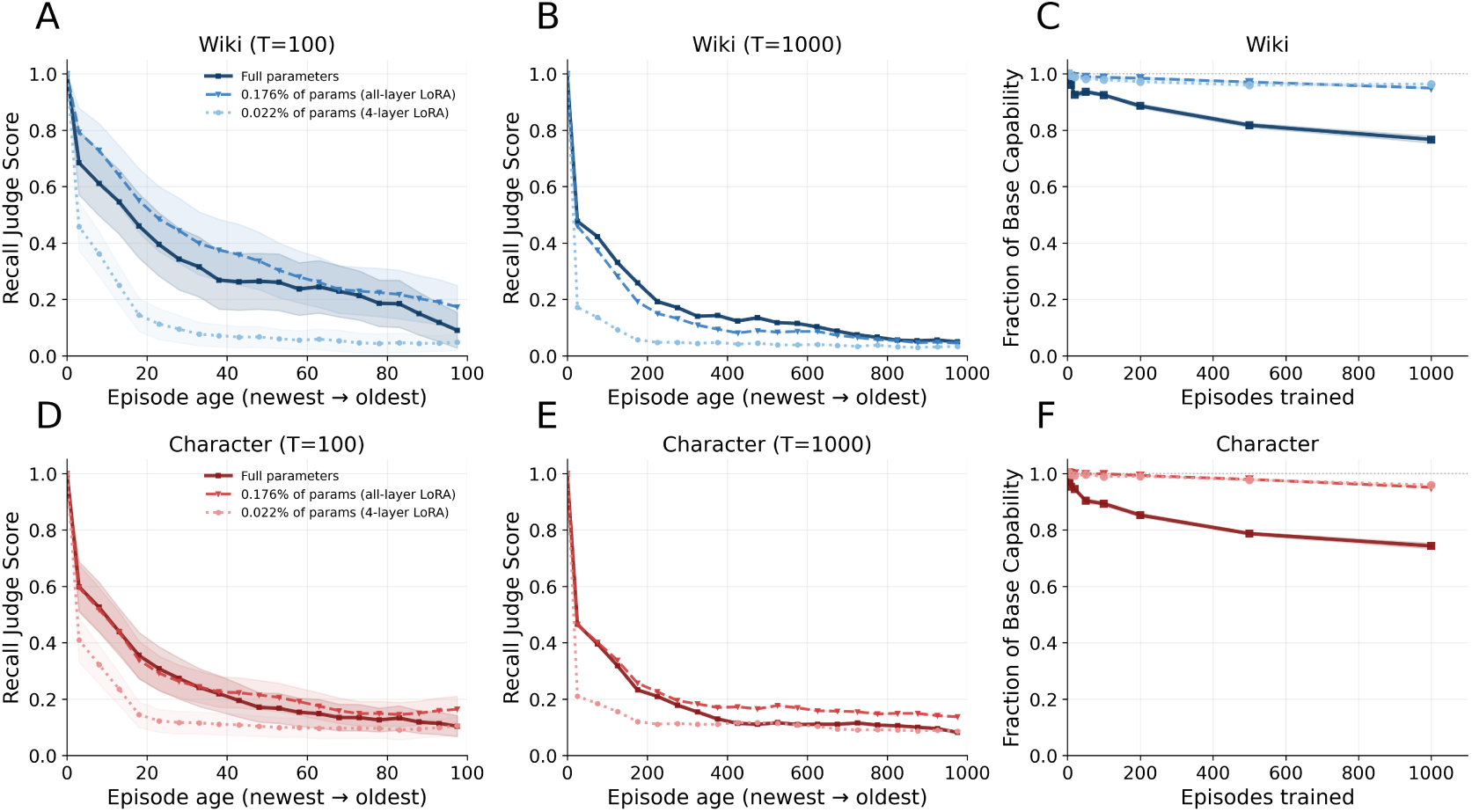
Catastrophic forgetting and capability degradation under continual fine-tuning. (A, B) Recall judge score as a function of episode age for the Wiki dataset after training on *T* = 100 (10 seeds for selecting different subsets of the dataset; shaded regions denote ±1 SEM) and *T* = 1,000 (1 seed, i.e. the full dataset) sequential episodes. (D, E) Same for the Character dataset. (C, F) Degradation of general capabilities (WinoGrande, HellaSwag, MMLU), measured as the fraction of the base model’s score retained as a function of the number of episodes trained. Full-parameter training degrades substantially after hundreds of episodes, while LoRA-based training largely preserves base capabilities on both datasets. Results shown are Llama; Qwen3-8B results in Supplementary Fig. 5.

### Regularization does not mitigate forgetting

Regularization is a common method for mitigating catastrophic forgetting in continual learning by constraining weight updates in a way that has minimal interference with existing memories. We evaluate three representative approaches: L2 regularization which penalizes deviation of the weights from their initial values, elastic weight consolidation (EWC) which penalizes changes to parameters deemed important for previously learned episodes (***Kirkpatrick et al., 2017***), and on-policy distillation (OPD) which regularizes the model’s output distribution to stay close to its predictions before each update (***Hinton et al., 2015***; ***Li and Hoiem, 2018***). Details of the implemented methods are in the Methods. For each method, we sweep the regularization strength over several orders of magnitude. The results reveal a consistent trade-off across all three methods (Fig. 2): weak regularization has a marginal effect on mitigating forgetting, while strong regularization prevents the model from encoding new episodes. No method finds an intermediate regime that preserves old episodes without sacrificing new ones. EWC is the most promising of the three, achieving the most uniform recall at intermediate strength on Wiki, but all episodes plateau around 50%, far from faithful retention.

**Figure 2.**
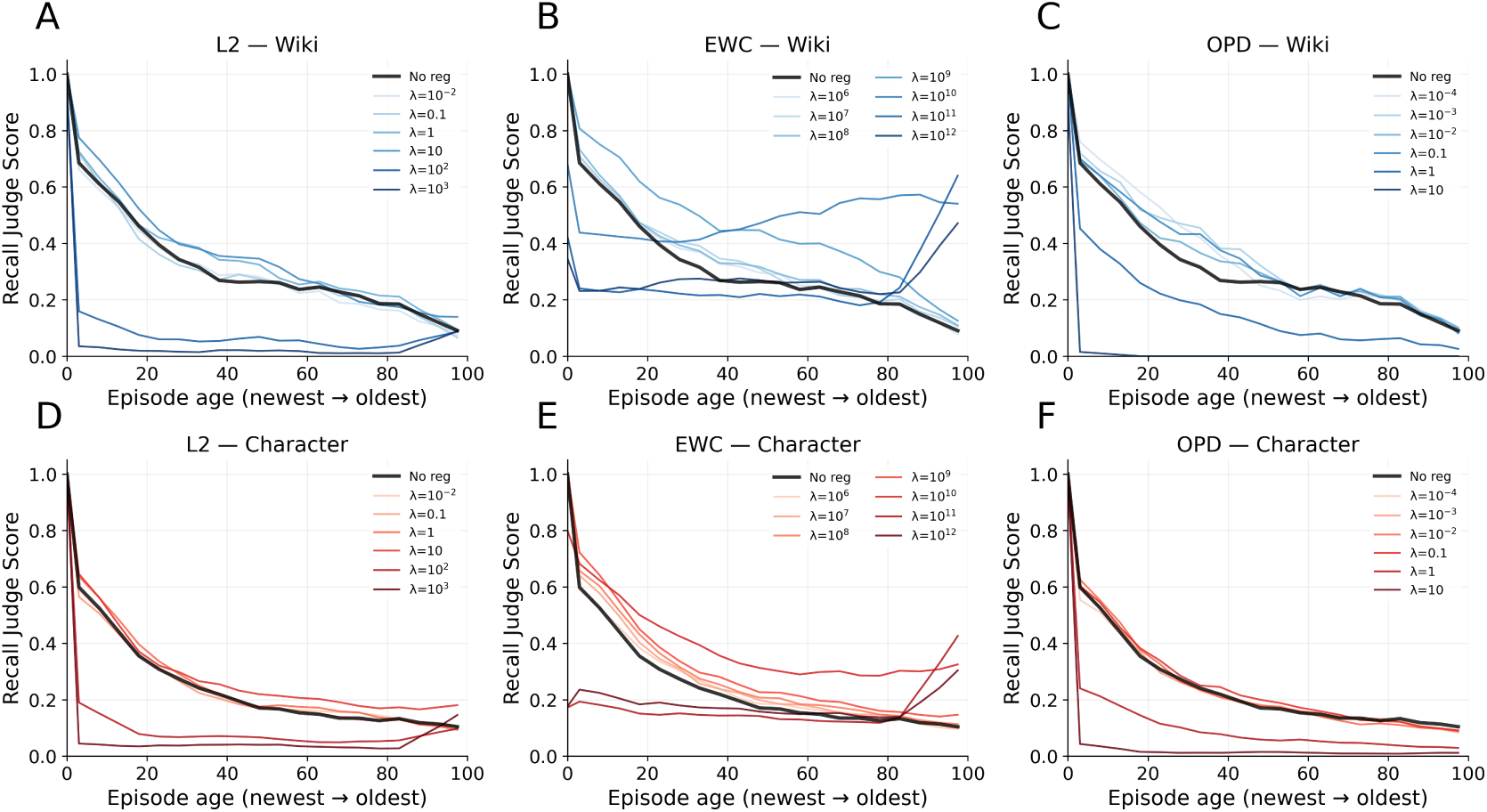
Effect of regularization on catastrophic forgetting under continual fine-tuning on 100 sequential episodes. Recall judge score is plotted against episode age (newest to oldest). (A, D) L2 regularization; (B, E) Elastic weight consolidation (EWC); (C, F) On-policy distillation (OPD). Top row: Wiki; bottom row: Character. Color gradient indicates regularization strength; black line shows the unregularized baseline. Across all three methods, weak regularization has no effect on the forgetting curve, while strong regularization suppresses learning entirely. Results shown are Llama; Qwen3-8B results in Supplementary Fig. 6.

### Complementary learning system for LLMs

As regularization did not solve the forgetting problem, we propose a memory system built from gated LoRA (***Wu et al., 2024***; ***Liang et al., 2026***), a computational analog of the brain’s complementary learning system (***McClelland et al., 1995***; ***Kumaran et al., 2016***) (Fig. 3). The pretrained base weights serve as a functional analog of the neocortex, holding stable knowledge; each low-rank adapter serves as a functional analog of the hippocampal engram that stores one episode; and the gating system is a winner-take-all competition that, at recall, activates only the most appropriate adapter.

**Figure 3.**
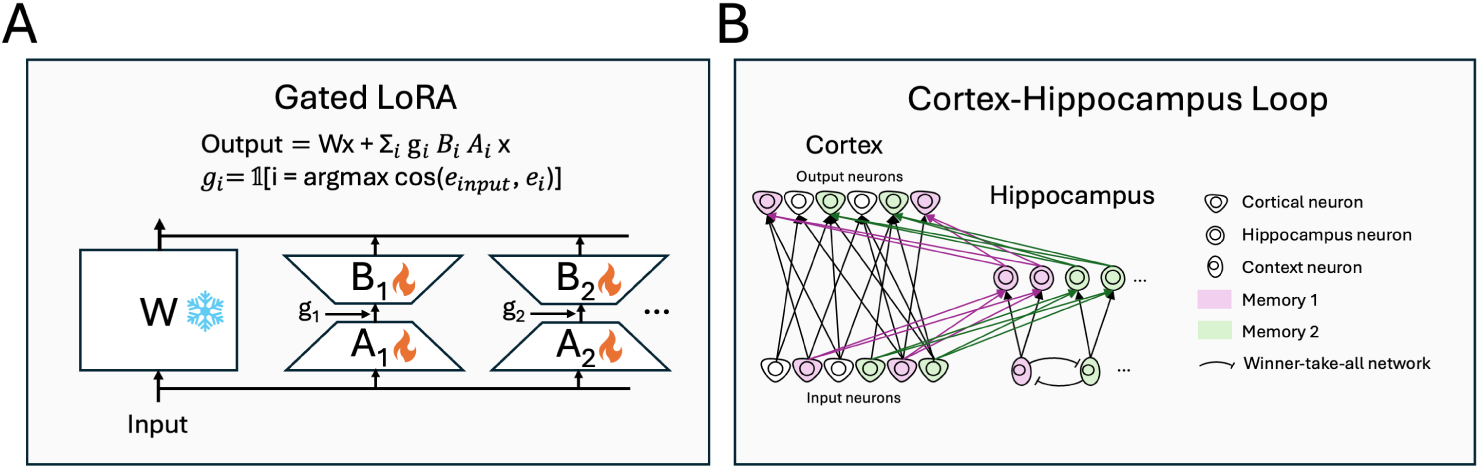
The gated LoRA and its correspondence to the cortex-hippocampus loop. (A) Gated LoRA. A frozen base weight matrix *W* is augmented with a set of per-episode low-rank adapters *B_i_A_i_*, each of which is gated by a gating signal *g_i_*. Only the adapter whose stored embedding best matches the input is activated. (B) The cortex-hippocampus loop has structural correspondence to gated LoRA. The frozen base weights play the role of cortex, holding stable long-term knowledge, while each adapter plays the role of a hippocampal trace that connects a sparse set of cortical neurons to a few hippocampus neurons and rapidly encodes one memory in a pattern-separated representation. Context neurons select the matching memory through a winner-take-all network, paralleling the embedding-similarity gating in (A), and hippocampal traces are gradually consolidated back into cortex.

Concretely, when a new episode arrives it is encoded into a fresh adapter on a small subset of the base model’s weight matrices; only this adapter’s parameters are trained, while the base model and all previously stored adapters are frozen by the gating. At inference, based on an incoming cue (a theme, a question, or partial information about an episode), the gating winner-take-all circuit chooses to open only the LoRA weights corresponding to the episode that best matches this cue. To implement this selection, during storage, the system computes an episode embedding vector, acting as a compressed summary of the episode. This vector is stored in the feedforward weights of the corresponding gating units. Similarly, during inference, the embedding of the cue is computed and input into all gating units, so that the system essentially computes the best match of the cue and stored episode embeddings. Thus, the output of the adapted layer becomes *W x* + ∑*_i_ g_i_B_i_A_i_x*, with the gate *g_i_* = *δ_i_*_,*i*max_ with *i*_max_ = arg max*_j_* cos(*e_j_*, *e*_cue_), where *e_j_* is the stored embedding of episode *j* and *e*_cue_ the embedding of the incoming cue. In practice, we use embeddings computed by a separate lightweight encoder (i.e., Qwen3-Embedding-0.6B (***Zhang et al., 2025***)); each embedding is stored alongside the adapter as its index. Since each episode is stored in a separate adapter, the total parameter count grows linearly with the number of episodes; parameter efficiency is therefore crucial to support a large number of stored episodes, which we investigate in the next section.

### Efficient single-episode storage

We ask how few parameters are needed to perfectly memorize a single episode. Standard LoRA already reduces the update to a low-rank factorization. To compress further, we take a rank-2 LoRA adapter on a single weight matrix and restrict the update to a randomly selected subset of *m* input and output neurons, so that only a subset of “cortical” neurons are connected to the “hippocampus”. We call this approach *masked* LoRA. By sweeping the mask size *m* across layers and weight types, we identify the minimal parameter count and its optimal placement that are sufficient for faithful recall. To quantify capacity, we take Wikipedia passages from a held-out set (distinct from the 1,000-article benchmark) and, for each one, binary-search over the mask size *m* to find the minimal number of trainable parameters (4*m* total, from two rank-2 matrices each with *m* active rows and columns) needed for perfect recall. We define memory capacity as the number of tokens stored per trainable parameter.

Figure 4A shows that capacity varies across layers. Middle layers (7–13) achieve the highest capacity across all weight types, peaking at approximately 0.5 tokens per parameter for the MLP down-projection at layer 10. Among weight types the MLP down-projection is the most efficient. The advantage of the middle layers is consistent with prior work localizing factual knowledge in mid-layer MLPs (***Geva et al., 2021***; ***Meng et al., 2022***; ***Dai et al., 2022***; ***Meng et al., 2023***). For the best configuration (layer 10, MLP down-projection), the minimal number of parameters scales linearly with passage length (Fig. 4B; *R*^2^ = 0.88), with a slope of approximately 2 parameters per token. At this rate, 1,000 passages of median length 76 tokens would require adapter parameters roughly 0.002% of the base model parameters. Qwen3-8B shows the same capacity profile over depth, the same ordering across weight types, and the same linear scaling of parameters with passage length, but its peak appears in slightly deeper layers: layer 15 of 36 at 0.48 tokens per parameter, against layer 10 of 32 at 0.51 in Llama (Supplementary Fig. 7). Adopting this efficient single-episode fine-tuning strategy, we proceed to evaluate the performance of the continually stored memories.

**Figure 4.**
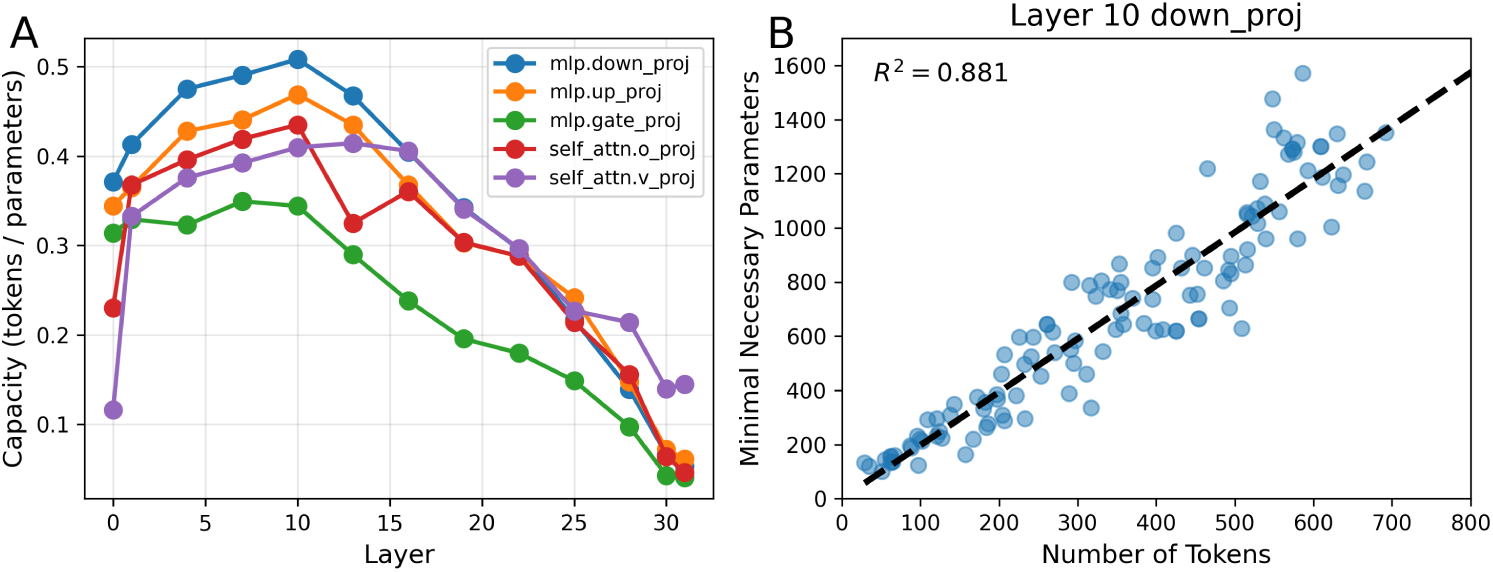
Parameter efficiency of masked LoRA for single-episode memorization in Llama. (A) Memory capacity (tokens stored per trainable parameter) across layers and weight types (the gate, up, and down projections of the MLP and the value and output projections of attention). The query and key projection of the attention layers failed to converge, thus they are omitted from the plot. Middle layers (7–13) achieve the highest capacity, with the MLP down-projection peaking at ∼0.5 tokens per parameter at layer 10. (B) Minimal number of trainable parameters versus passage length for the best configuration (layer 10, MLP down-projection; *n* = 121 passages). The linear fit (*R*^2^ = 0.88) yields a slope of ∼2 parameters per token. Results shown are Llama; Qwen3-8B results in Supplementary Fig. 7.

### Storing and retrieving a thousand episodes

The gated LoRA framework is implemented, together with the efficient fine-tuning of single episodes, as follows. For each new episode, a fresh masked LoRA adapter of rank 2 is attached to the MLP down-projection at layer 10, identified as the most parameter-efficient location in the preceding section. The initial mask size is set at approximately 2.5 parameters per token (*m* = 45 for Wiki and *m* = 60 for Character), following the linear scaling in Fig. 4B. If the initial mask is insufficient for perfect reconstruction, an auto-growth mechanism expands the mask by a factor of 1.2 and retrains until exact recall is achieved. In practice, the resulting set of adapters is extremely compact: across 1,000 Wiki passages, total adapter parameters sum to 197K (0.0025% of the base model, 2.6 parameters per token), while 1,000 Character episodes require 256K parameters (0.0032% of the base model, 2.7 parameters per token). All adapters are trained with the same prompt (“Reconstruct the story:”), so the adapter weights alone determine which passage is reconstructed.

Unlike the setup in the section *Failures of continual fine-tuning*, where the model is trained to map a theme cue to the passage, we first test whether the model can use a question related to one of the episodes as a cue to recall the episode. On the Wiki dataset, question-cued recall remains near-perfect across the entire range, declining only from 100% to 99.5% at *N* = 1,000 (Fig. 5). The Character dataset shows a more noticeable decline, from 100% at small *N* to 86.8% at *N* = 1,000. The decline is driven by errors in embedding-based routing: when the correct adapter is selected, reconstruction is near-perfect regardless of dataset. The greater drop on the Character dataset reflects the tighter semantic overlap among a single individual’s life narratives compared to topically diverse Wikipedia articles. We additionally show routing performance with theme cues in Supplementary Fig. 3. Because the short theme labels carry less semantic information than a full question, and because the fictional character’s episodes share recurring themes, theme-cued routing degrades more steeply with scale: at *N* = 1,000, top-1 theme routing accuracy falls to 65.6% on the Character dataset (versus 96.5% on Wiki).

**Figure 5.**
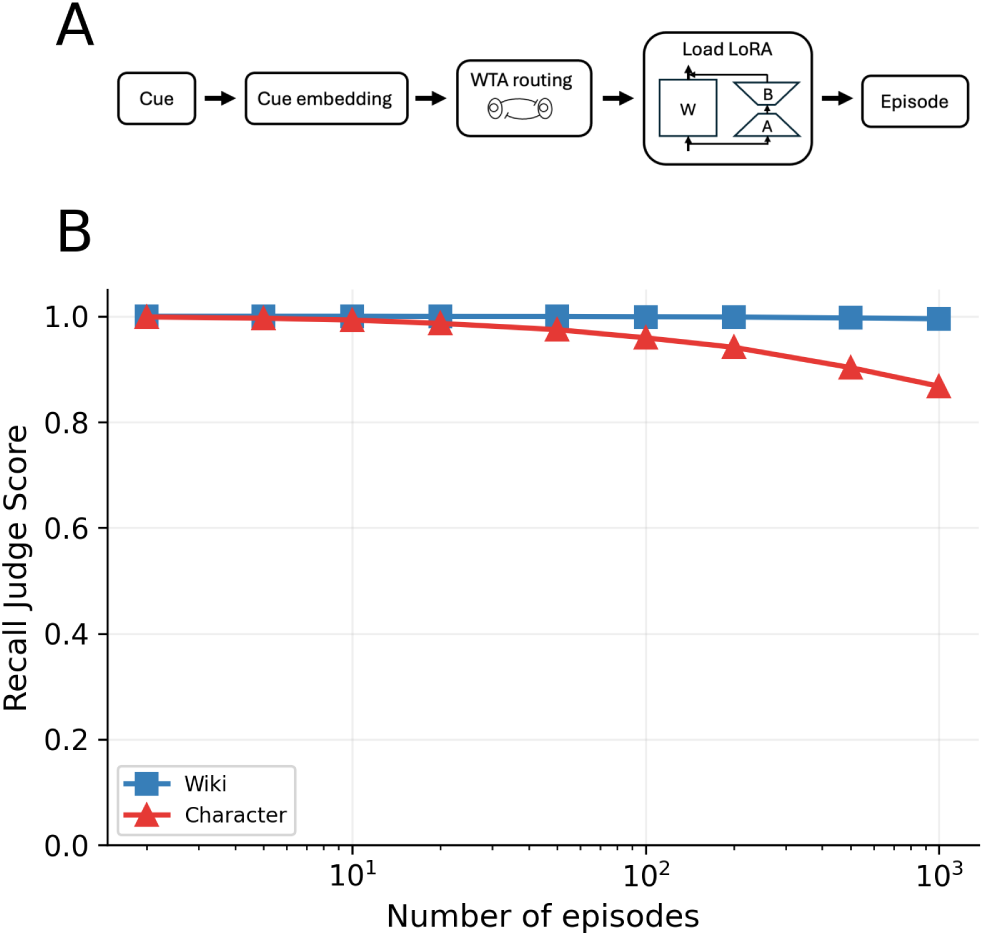
End-to-end recall with the gated LoRA framework. (A) The recall pipeline: a cue is embedded, routed by winner-take-all selection to its best-matching adapter, which is loaded onto the base model to reconstruct the episode. (B) Recall judge score as a function of the number of stored episodes. Wiki passages (blue) exhibit very minimal forgetting, maintaining ∼99% recall at *N* = 1,000. Character episodes (red) show a gradual decline to 86.8% at *N* = 1,000, reflecting the greater difficulty of routing among semantically similar episodes. Values below *N* = 1,000 average 100 random subsets; *N* = 1,000 uses the full dataset and a single adapter-training run (seed 72). Results shown are Llama; Qwen3-8B results in Supplementary Fig. 8.

To understand this bottleneck, we evaluate the routing accuracy directly as a function of *N* (Fig. 6A). On the Wiki dataset, question-cued routing stays near-perfect across the full range. On the Character dataset, top-1 question-based routing drops to 85.2% at *N* = 1,000, which is slightly lower than the recall judge score in Fig. 5 due to the best distractor episode often containing some overlapping content with the target, but it recovers to 96.0% at top-5, where routing counts as correct if the target episode is among the five best-matching episodes for the cue. This gap reflects the fact that the embeddings cluster more tightly on the Character dataset. Its pairwise episode-episode cosine similarities are shifted markedly to higher values than those of Wiki (Fig. 6B). The consequence for routing is shown in Fig. 6C, which compares, across cues, the similarity of each cue to its correct episode against its similarity to the closest distractor episode. On Wiki these two distributions are well separated, so the correct episode almost always wins the match. On Character they overlap, as the correct-pair similarity is lower and the best-distractor similarity is higher.

**Figure 6.**
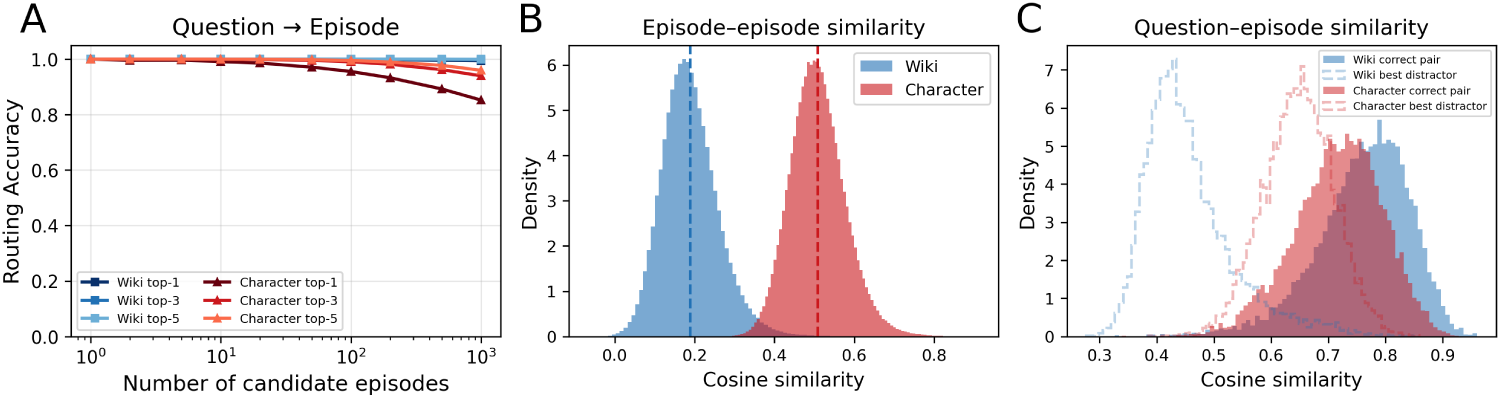
Embedding-based routing accuracy and similarity structure. (A) Question-to-episode routing accuracy (top-1, top-3, top-5) as a function of the number of candidate episodes; question embeddings provide a strong routing signal, reaching 99.4% top-1 on Wiki and 85.2% on Character at *N* = 1,000. Panel A is computed on the complete, unfiltered set of 10 questions per episode; values below *N* = 1,000 average 50 random subsets, whereas *N* = 1,000 uses the full dataset. (B) Distribution of pairwise episode-episode cosine similarities; Character embeddings cluster more tightly than Wiki, explaining the greater routing difficulty. (C) Distribution of question-episode cosine similarities for correct pairs versus best distractors; on Wiki the two distributions are well separated, while on Character they overlap substantially. Because the embedding encoder and datasets are common to both base LLMs, these routing analyses are model-independent; they are reproduced with the Qwen results in Supplementary Fig. 9.

### Question-answering ability via internal RAG

Faithful reconstruction is a measure of memorization. Here we seek to explore ways in which the fast hippocampus-like adapters can be flexibly used as knowledge to answer factual questions about stored episodes. We first tested routing the question to the best-matching adapter via embedding similarity, composing the adapter with the base model, and generating an answer directly. This direct method fails on both datasets (Fig. 7B). It indicates that the adapters trained to recall the episodes faithfully do not generalize to direct question-answering abilities. This failure is consistent with previous findings (***Ovadia et al., 2024***; ***Mecklenburg et al., 2024***; ***Zhao et al., 2026***; ***Lampinen et al., 2025***).

**Figure 7.**
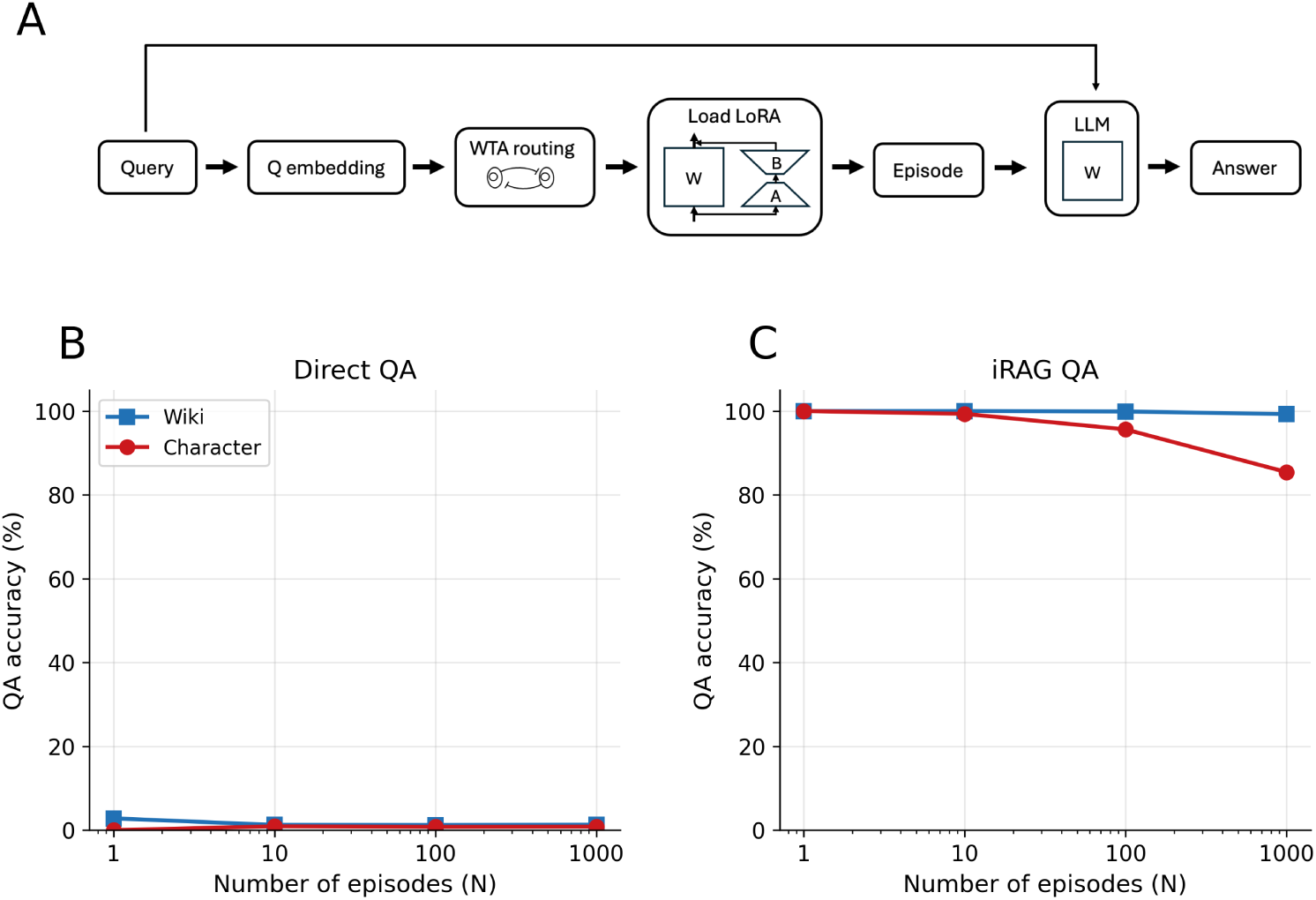
Question answering over stored episodes. (A) The iRAG pipeline. An incoming cue is embedded and routed by winner-take-all (WTA) embedding selection to its best-matching adapter, which is loaded onto the base model to reconstruct the stored episode. The reconstructed passage is then returned to the base model’s context, and the LLM answers the question over it. (B) Direct QA, where the question is answered by the adapter-augmented model without reconstruction, achieves near-zero accuracy (∼1%) on both datasets. (C) Under the filtered evaluation, iRAG achieves 99.3% on the Wiki dataset and 85.4% on the Character dataset at *N* = 1,000. Accuracy in (B) and (C) is scored on the filtered question set defined in Methods, which removes questions the base model already answers and questions not answerable from the source passage in context; by construction this makes iRAG accuracy 100% at *N* = 1 in (C). Unfiltered accuracies and the counts removed by each criterion are given in Supplementary Note 2. Results shown are Llama; Qwen3-8B results in Supplementary Fig. 10. Confidence intervals over episodes are below ±1.5 percentage points and are omitted for clarity (Methods).

To bridge this gap, we introduce *internal RAG* (iRAG), a two-stage pipeline inspired by Retrieval-Augmented Generation (***Lewis et al., 2020***; ***Brown et al., 2020***) that leverages both the adapter’s reconstruction ability and the base model’s in-context learning capability. In the first stage, the incoming question is routed to the best-matching adapter via embedding similarity to recall the stored episode. In the second stage, the adapter is removed, and the reconstructed passage is placed in the base model’s context window followed by the question. The base model then answers the question using standard in-context QA over the reconstruction. Such a two-stage process resembles retrieving episodic memories into working memory for reasoning in the human memory system. Leveraging iRAG, the accuracy implied by top-1 routing is ∼99% on the Wiki dataset at *N* = 1,000 and ∼85% on the Character dataset.

Note that, in the main experiments, we filter out questions that can be answered by either base model without the episode in context, or that either base model fails to answer even with the episode in context (Methods). The former removes questions whose answers are already accessible to either base model under the tested prompting conditions, while the latter ensures that the question is actually answerable and can be correctly judged. Both criteria are applied for Llama-3.1-8B-Instruct and then again for Qwen3-8B, leaving 5,873 of 9,982 Wiki questions and 5,211 of 10,000 Character questions. Because of this dataset design, iRAG tests whether the system routes a question to the correct episodic trace and reconstructs that episode into context. Conditional on correct routing and reconstruction, the base model answers these questions successfully by construction. The decrease in the number of stored episodes therefore measures interference at the routing stage.

### Transferring memories to base model: reconstruction

We model gated LoRA adapters as a fast-learning, hippocampus-like memory system. Over time, their contents are consolidated into the base model weights, providing a cortex-like long-term store and allowing adapter capacity to be freed for new experiences. This process parallels the replay-based consolidation thought to occur in the mammalian brain (***Wilson and McNaughton, 1994***; ***McClelland et al., 1995***). Here we mimic the effect of replay by fine-tuning the base model parameters to memorize a set of episodes in batch mode. Each fine-tuning step consists of small mini-batches, resembling the consolidation by replay of a group of stored episodes. This base-weight fine-tuning is implemented as a high-rank LoRA update (i.e. rank 2048) that is merged back into the base weights after training and then removed (see Methods). Since in the batch mode, the same set of parameters stores all the memories, we need to determine which specific prompts need to be added to each episode during the fine-tuning such that during recall a corresponding recalling cue would trigger the reconstruction of the relevant episode. We first use a cue that contains the episode’s embedding vector. Specifically, given the prompt “Reconstruct the story about <EMB=”, where <EMB= is a placeholder token whose input embedding is replaced by the stored episode embedding, the model is trained to generate the corresponding passage. This design enables the base model weights to achieve the same goal as LoRA adapters, i.e. retrieving the passage that corresponds to an episode embedding. On both datasets, this method achieves near-perfect recall (Fig. 8B), demonstrating that the consolidated model can reconstruct all 1,000 episodes from their embeddings. We have also demonstrated an excellent reconstruction using natural language cues in the form of the episodic themes, Supplementary Fig. 2A.

**Figure 8.**
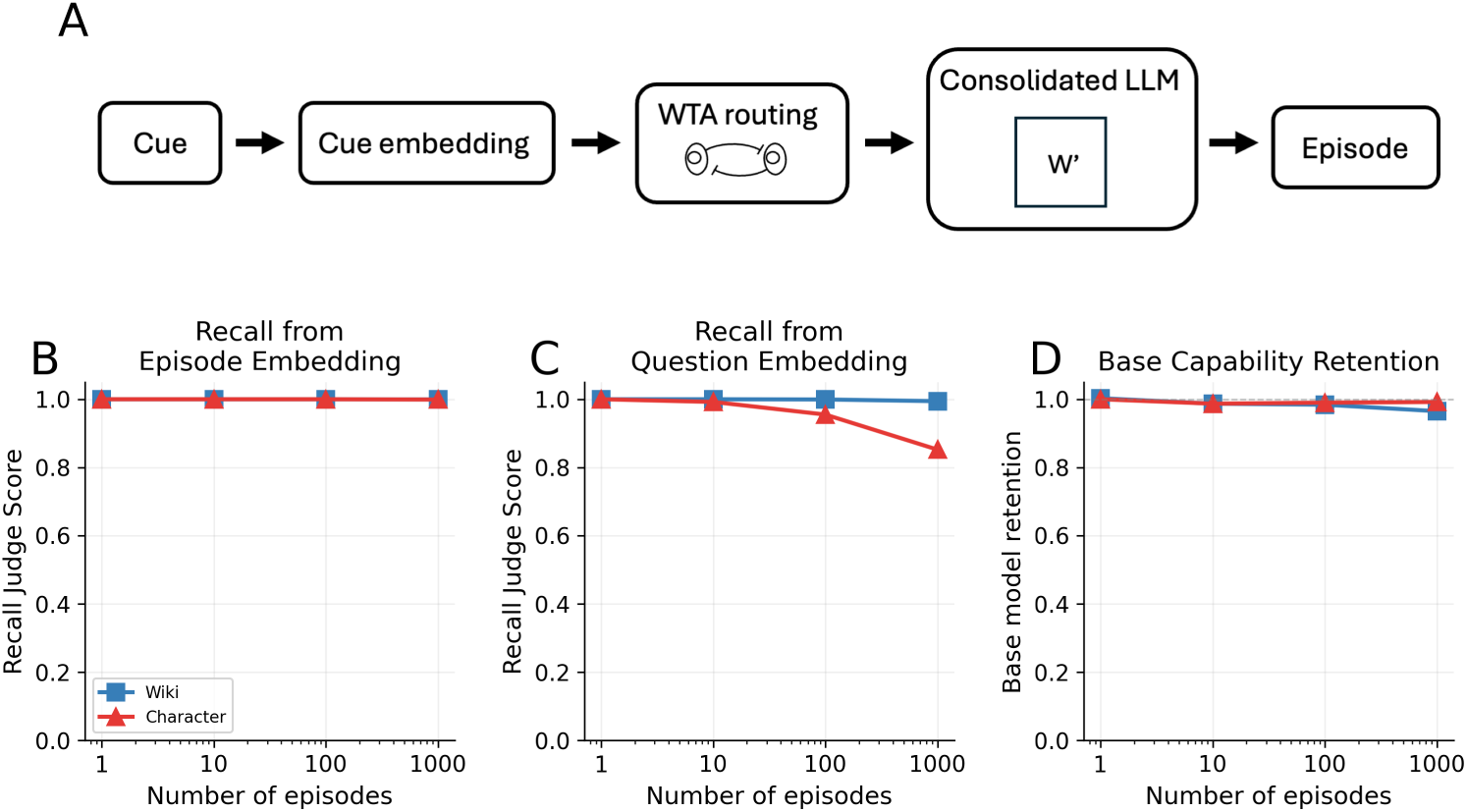
Embedding-cued reconstruction after consolidating the stored episodes into the base model’s weights. (A) Embedding-cued recall with the consolidated model. A cue is embedded and routed by winner-take-all (WTA) similarity to its nearest stored episode embedding, which is fed to the consolidated model (weights *W* ^′^) to reconstruct the episode, with no adapter loaded. (B) Recall when the model is cued directly with the correct episode embedding; near-perfect across all numbers of stored episodes and both datasets. (C) End-to-end recall with a *routed question-embedding cue*: the question embedding is routed to its nearest stored episode embedding, which then cues reconstruction. This degrades only on the Character dataset at large *N*, reflecting the embedding overlap that limits routing accuracy (Fig. 6); it is distinct from the direct question-embedding cue in Supplementary Fig. 2B. (D) Fraction of base model capability retained after consolidation, measured by average performance on WinoGrande, HellaSwag, and MMLU. Each *N* is a single consolidation run; *N* = 1,000 uses the full dataset. Results shown are Llama; Qwen3-8B results in Supplementary Fig. 11.

On the other hand, when the model is consolidated with cues of episodic theme, triggering recall by cues of relevant questions (as a text prompt) suffers substantial degradation over 1, 000 stories, particularly for the Character dataset (see Supplementary Fig. 2A), indicating that in contrast to themes, which are appended to each story during consolidation, a typical question is only partially successful in disentangling the relevant story from the cumulative memory traces. When the model is consolidated with cues of episodic embedding, triggering recall by prompting on the question embedding was successful for the wiki data but suffered a substantial loss in the character dataset (see Supplementary Fig. 2B). To improve the retrieval of the relevant passages based on a question, we employ an explicit routing system similar to the iRAG above. Here, a network stores all the memorized episodes’ embeddings in the feedforward weights of a WTA circuit, and this circuit selects the one that best matches the embedding of an incoming question. Specifically, during recall, we first compute the embedding of an incoming question, which is used in the WTA circuit to select the best-matching embedding. Second, the winning episodic embedding is input into the context window to trigger recall of the episode with the matching embedding. This *routed question-embedding cue* closely tracks iRAG (Fig. 8C). Crucially, this consolidation leaves the base model’s general capabilities largely intact across all numbers of stored episodes, retaining 94–95% of the base score without augmentation (Fig. 8D).

### Transferring memories to base model: QA

We next ask how the consolidated memories can be used as new knowledge, as measured by answering questions related to the memorized episodes without recalling the passage. We first test whether the consolidated model introduced in the previous section can answer questions directly from its parameters, without first reconstructing the passage. Direct QA works poorly on the model fine-tuned to reproduce the original passages, 31%/33% (Wiki/Character) (Fig. 9C). Such generalization problem of knowledge injection by fine-tuning is well known (***Ovadia et al., 2024***; ***Mecklenburg et al., 2024***). The failure is not solely due to interference between 1,000 episodes; accuracies at *N* = 1 are low as well, 43%/51% (Wiki/Character) (Fig. 9B). The failure arises because the training objective of minimizing reconstruction loss on a single passage is insufficient to induce the model to use the acquired knowledge flexibly.

**Figure 9.**
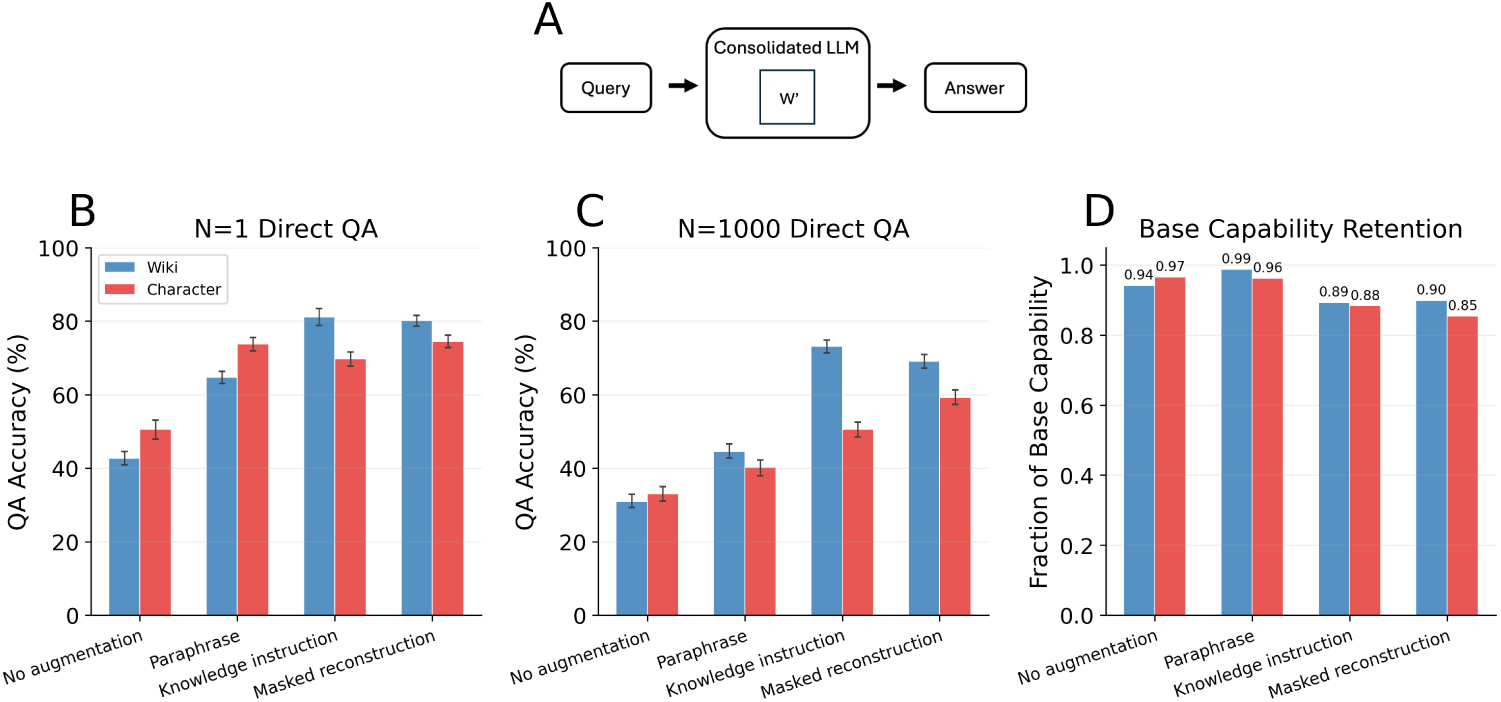
Direct QA after consolidation. (A) Direct QA with the consolidated model. A query is fed directly to the consolidated model (weights *W* ^′^), which answers without loading any adapter or reconstructing the passage. (B) Direct QA accuracy at *N*=1 under four training conditions (no augmentation, paraphrase, knowledge-instruction, and masked reconstruction), evaluated by LLM judge. (C) Direct QA accuracy at *N*=1,000 under the same four training conditions, evaluated by LLM judge. (D) Fraction of base model capability retained after consolidation at *N*=1,000, measured by average performance on WinoGrande, HellaSwag, and MMLU. Results shown are Llama; Qwen3-8B results in Supplementary Fig. 12.

On top of simply fine-tuning the model to reconstruct the stored passages, we tested three augmentation strategies of increasing aggressiveness (details in Methods). Paraphrase augmentation adds paraphrases of each passage as additional reconstruction targets (***Ovadia et al., 2024***). Building on paraphrase augmentation, knowledge-instruction augmentation adds QA-format declarative facts extracted from each passage (***Ovadia et al., 2025***), while masked reconstruction instead adds a denoising objective in which the model recovers a passage from a partially masked copy (Pan et al., 2025a). At *N* = 1, paraphrase augmentation raises accuracy to 65%/74% (Wiki/Character), knowledge-instruction augmentation to 81%/70%, and masked reconstruction to 80%/74% (Fig. 9B). At *N* = 1,000, the corresponding accuracies are 45%/40%, 73%/50%, and 69%/59% (Fig. 9C). Since even a single episode yields accuracy largely below perfect, the primary bottleneck appears to be the knowledge injection procedure itself rather than inter-episode interference. Nevertheless, the drop from *N* = 1 to *N* = 1,000 is consistently larger on the Character dataset than on Wiki across all conditions, suggesting that interference between semantically correlated episodes still makes knowledge internalization harder. The gains in direct QA accuracy come at a modest cost to the model’s general capabilities. Relative to the two milder conditions, the more aggressive knowledge-instruction and masked-reconstruction augmentations generally yield higher direct QA accuracy with lower base-capability retention (Fig. 9C,D); the relationship is not strictly monotone, particularly for paraphrase augmentation on Wiki. However, this trade-off is relatively mild: even the most damaging augmentation, masked reconstruction, retains 90%/85% (Wiki/Character) of the base model’s average performance on WinoGrande, HellaSwag, and MMLU.

These results indicate that fully internalizing episodic memories into flexible parametric knowledge remains a significant challenge, though augmentation during consolidation narrows the gap.

## Discussion

### A functional model of continual episodic memory

Our results show that the principal operations associated with complementary learning systems can be implemented in an LLM: rapid storage in separated episode-specific traces, competitive retrieval from a cue, and slower consolidation into shared model weights. Separating the traces prevents catastrophic interference during storage, while semantic routing allows the appropriate trace to be selected during recall.Consolidation preserves cue-dependent reconstruction without requiring an episode-specific adapter, while also making the stored content partially accessible for direct question answering. Together, these mechanisms provide a concrete neural-network instantiation of a hippocampus–cortex division of labor.

### Pattern separation through private traces

Our design is motivated by pattern separation in the hippocampus, where sparse coding is thought to reduce overlap among correlated memories (***O’Reilly and McClelland, 1994***; ***Yassa and Stark, 2011***). There are at least two mechanistically distinct ways to obtain separated traces. In sparse expansion, a dense representation is projected into a much larger space and thresholded, allowing a fixed transformation to decorrelate different inputs (***Babadi and Sompolinsky, 2014***). In engram allocation, a small population of neurons is recruited to store each new memory (***Guan et al., 2016***; ***Tonegawa et al., 2015***; ***Josselyn and Tonegawa, 2020***).

Our framework is closer to the engram view. Each episode is assigned its own private adapter, so different traces have zero parameter overlap by construction. Sparsity determines the resource cost of each adapter rather than producing the separation itself. This differs from biological engram formation, in which neurons compete for allocation to a memory trace and more excitable neurons are preferentially recruited, for example through CREB-dependent mechanisms (***Stefanelli et al., 2016***; ***Rao-Ruiz et al., 2019***; ***Kim et al., 2013***; ***Han et al., 2007***; ***Josselyn et al., 2015***). In the present model, allocation during storage is predetermined: each incoming episode receives a fresh adapter. The model therefore captures the functional consequence of engram allocation, namely separated traces, but not the competitive process through which biological traces are formed.

Biological engrams are also dynamic. Their neuronal composition and tuning can drift over time (***Rule et al., 2019***), and memories formed close together may occupy partially overlapping ensembles that link the corresponding experiences (***Cai et al., 2016***). Such overlap may support integration, generalization, and memory updating (***Tonegawa et al., 2015***; ***Josselyn et al., 2015***). By contrast, the present adapters are static and nonoverlapping. Allowing adapters to merge, split, overlap, or reorganize according to semantic similarity could make more efficient use of limited memory capacity and provide a model of interactions among related experiences.

### Retrieval mechanism

Pattern separation keeps memory traces distinct, but recall must still select and reconstruct the appropriate memory from a cue. In classical autoassociative memories, retrieval is defined as pattern completion: the cue is a partial or corrupted version of a stored pattern, and recurrent attractor dynamics recover the complete pattern (***Hopfield, 1982***; ***Kanter and Sompolinsky, 1987***; ***Treves and Rolls, 1994***). For contextualized, semantically rich episodes, however, there is no equally natural definition of a partial pattern. A theme, a question, a partial description, and the beginning of an episode provide qualitatively different routes to the same memory. This ambiguity motivates memory architectures that separate the address used for retrieval from the content being retrieved. Scaffold-based content-addressable memories use a fixed scaffold for addressing and separate weights for storing content (***Sharma et al., 2022***; ***Chandra et al., 2025***). Hippocampal indexing theory similarly proposes that the hippocampus stores an index to a memory whose content is distributed across the neocortex; reactivating the index reinstates the associated cortical pattern (***Teyler and Rudy, 2007***). This organization can also be expressed as a key-value memory (***Gershman et al., 2025***).

In our model, the semantic embedding of an episode serves as its retrieval key, while the corresponding adapter stores the episode’s content. In the experimental implementation, the cue embedding is compared with the stored episode embeddings, and the adapter with the largest cosine similarity is selected. The same computation can be implemented by a winner-take-all neural circuit. Each episode embedding is stored in the feedforward afferent weights of one gating unit, which compute the dot product between the episode embedding and the incoming cue embedding. Recurrent lateral inhibition among the gating units then implements the competition that selects the unit receiving the strongest similarity signal. The feedforward weights therefore compute semantic similarity, while recurrent competitive dynamics perform selection.

This organization gives the address a defined semantic content and specifies how heterogeneous cues are connected to the appropriate memory trace. It also separates routing from reconstruction: the embedding identifies the relevant episode, whereas the selected adapter reconstructs its passage.

### From episodic recall to usable knowledge

Our results separate two problems that are often treated together: storing new information and making that information usable as knowledge. Gated LoRA supports highly accurate recall of stored episodes, but directly answering a question from the selected adapter is substantially more difficult. This agrees with previous findings that fine-tuning on new text can teach a model to reproduce the text without making its contents reliably available for downstream questions and reasoning (***Ovadia et al., 2024***; ***Mecklenburg et al., 2024***; ***Lampinen et al., 2025***).

Knowledge editing addresses a related but narrower problem. ROME and MEMIT modify selected weights to introduce atomic factual associations (***Meng et al., 2022***, ***2023***), whereas MELO and WISE isolate edits in routed parameter modules (***Yu et al., 2024***; ***Wang et al., 2024***). MELO is particularly close to our architecture because it uses similarity-based routing to activate LoRA parameters. Its evaluation, however, primarily asks whether paraphrases of edited factual questions produce the intended short answers. Our task requires storing and reconstructing a semantically rich episode, then extracting information from it in response to a new question. The expected answer need not reproduce the wording of either the passage or the training questions.

WISE similarly uses a routed side memory to reduce interference among sequential factual edits (***Wang et al., 2024***). Success on factual-editing benchmarks does not establish that an extended episode can be reconstructed through free generation or that its contents can be flexibly used in a broader context. The present results therefore expose a distinction that is less visible in factual editing: accurate storage of an episode does not guarantee direct access to the knowledge contained in it.

Training with additional objectives improves this access. Paraphrases (***Ovadia et al., 2024***), question-answer pairs extracted from stored passages (***Ovadia et al., 2025***), and demasking objectives (Pan et al., 2025a) all increase direct question-answering accuracy. None eliminates the advantage of first reconstructing the relevant episode in context. This explains the effectiveness of iRAG: reconstruction makes the episode immediately available to the language and reasoning mechanisms already present in the base model. A complementary direction would be to encode each episode together with representations that explicitly relate it to the model’s existing knowledge.

Batch consolidation exposes a related limitation in reconstruction. Theme cues can faithfully reconstruct the episodes from the shared weights, demonstrating that the individual memories remain encoded, whereas questions are substantially less effective at directly triggering their reconstruction. It remains unclear whether the limitations of question-cued reconstruction and direct QA arise primarily from the autoregressive training objectives of current LLMs, and therefore may not generalize to human memory, or reflect a more general challenge of retrieving and using semantically structured memories encoded in distributed representations.

### Relation to episodic-memory and consolidation architectures

Existing memory architectures for LLMs differ in what they store and in the role memory serves. ***Spens and Burgess (2026)*** similarly model hippocampo-neocortical interaction using an LLM, but focus on the compressed representation of episodes in hippocampal memory and the ability of generative semantic memory to fill in predictable information during reconstruction. RAG provides the interface between the two systems: a cue retrieves a compressed hippocampal trace, consisting of a gist vector and unpredictable details, which conditions the LLM’s reconstruction. However, they do not evaluate the hippocampal store under continual addition of episodes, and they explicitly leave hippocampal forgetting and retrieval failures outside the model. Consequently, they do not measure interference or forgetting curves in fast episodic memory. They also do not evaluate factual QA through the hippocampal RAG pathway. Direct factual QA is tested only after consolidation into the LLM, without RAG, using a three-alternative embedding-similarity criterion. In contrast, the present work focuses on three related challenges. The first is preventing interference as episodes are encoded continually. The second is storage efficiency: fast episodic memory must allocate a distinct trace to each incoming episode while operating with finite resources. The third is transforming memorized episodes into knowledge accessible through QA, both in fast memory and after consolidation. Using the LLM as a judge that measures the recovery of the queried core fact, we show that transforming memorized episodes into knowledge accessible through QA remains a substantial challenge.

HippoRAG constructs an external knowledge graph and retrieves relational evidence for multihop question answering (***Gutiérrez et al., 2024***). Larimar stores factual prompt-answer associations in an external associative-memory matrix that supports rapid updating and selective forgetting (***Das et al., 2024***). These systems provide effective access to externally stored information, but they do not encode rich episodes as distinct parametric traces or model their subsequent integration into the base network.

“Language Models Need Sleep” addresses consolidation using a hierarchy of memories, distillation, parameter expansion, and reinforcement learning over self-generated rehearsal data (***Behrouz et al., 2026***). Our results show that direct batch replay of the original passages provides a simpler route from episode-specific traces to shared model parameters. Replay preserves cue-dependent reconstruction, but it does not fully transform episodic content into flexibly accessible semantic knowledge. Determining what should be replayed, and whether consolidation should preserve individual episodes or extract relational and semantic structure from them, remains an open problem.

Broader accounts of human-like episodic memory include additional processes such as event segmentation, selective encoding, temporal organization, memory updating, competitive retrieval, and consolidation into semantic knowledge (***Dong et al., 2025***). Related proposals for long-term LLM agents connect rapid episodic storage, reinstatement into active context, and gradual incorporation into model parameters (***Pink et al., 2025***). The present model provides a concrete implementation of separated parametric traces, competitive semantic retrieval, and replay-based consolidation for rich textual experiences.

### Limitations and future directions

Our study leaves several questions open. First, consolidation is performed offline after all episodes have been encoded. A lifelong system would need to interleave encoding and consolidation, determine which memories to replay, and decide when an adapter can be discarded to free capacity. Scheduling these operations has been studied in continual learning (***Horgan et al., 2018***; ***Rolnick et al., 2019***; ***Klasson et al., 2023***), but not in an LLM that combines episode-specific traces with replay-based consolidation. A biologically motivated possibility is spontaneous replay, in which noise in the gating circuit occasionally reactivates an adapter and causes its episode to be replayed. In related attractor models, noise-driven reactivation can occur at a rate determined by a memory’s basin of attraction (***Shaham et al., 2022***); in the present setting, replay frequency might instead depend on salience, recency, or retrieval history.

Second, routing relies on a separate general-purpose embedding model and becomes the main source of error for the correlated Character episodes. Contrastive training, learned indexing, or routing based on the memory model’s own representations could improve retrieval and make the index more fully internal to the system.

Third, it remains unknown whether one embedding and one adapter per episode will scale from 1,000 memories to tens or hundreds of thousands of memories. At larger scales, efficient retrieval and storage may require hierarchical organization, shared structure among related memories, or mechanisms that merge and split traces over time.

Fourth, the datasets consist of clean, pre-segmented episodes, and the tasks do not require the model to represent temporal relations among episodes. Human experience instead arrives as a continuous stream that is spontaneously divided into events, with event boundaries reflected in behavior and hippocampal activity (***Zacks et al., 2007***; ***Baldassano et al., 2017***). Extending the model to discover episode boundaries and represent temporal relations would be necessary for a fuller account of continual experience.

Finally, the present study evaluates memory primarily against faithful reconstruction and factual question answering. Human memory is also lossy and reconstructive. Forgetting follows regular temporal patterns (***Wixted and Ebbesen, 1991***), and recall can produce schema-consistent distortions and confident false memories (***Roediger and McDermott, 1995***; ***Schacter et al., 2011***). Routing errors in the present system provide a simple form of memory substitution, but they do not yet reproduce the structured distortions characteristic of human recall. Modeling such errors, rather than only minimizing them, could make the framework useful for studying how episodic memories are transformed over time.

## Methods

### Base models

Every experiment is run with two base models, Llama-3.1-8B-Instruct (***Grattafiori et al., 2024***) and Qwen3-8B (***Yang et al., 2025***). The main figures report Llama; the matching Qwen figures are in Supplementary Note 7. The two sets of runs share the datasets, the training harness, the hyperparameters and the LLM judge. The only differences are architectural, together with the corresponding input templates. The QA filter defined below is applied against both base models in turn, so the same question set is used for both and their accuracies are directly comparable.

In both models, during the fast encoding of episodes into LoRA adapters, the base model’s weights remain frozen. During consolidation, the base weights are modified. For computational efficiency, we implement this base-weight update as a high-rank LoRA (i.e. rank 2048): we first train the model with the LoRA, then merge the LoRA weights into the base weights, so that at inference the modified weights carry the memories and no adapter is attached. LoRA here is therefore a compute-saving parameterization of a direct base-weight modification, not a separate persistent store.

### Datasets

#### Wikipedia dataset

We crawled Wikipedia passages released in 2026, well after the post-training knowledge cutoffs of both base models. Length filters (65–125 tokens under the Llama tokenizer) and diversity filters (title-overlap deduplication, exclusion of list-like entries) yielded 2,030 candidate articles, from which we sampled 1,000 with a fixed random seed. Passages were lightly cleaned by a language model to correct formatting and template artifacts while preserving factual content. For each passage, gpt-5.2-2025-12-11 generated 10 self-contained factual question– answer pairs (maximum 5 words per answer) and 9 paraphrases emphasizing word-, sentence-, and paragraph-level reordering. Final passage statistics: 76.8 ± 9.6 tokens (mean ± s.d.; median 76, interquartile range 69–85).

#### Character dataset

We constructed a synthetic autobiographical dataset for a single fictional character, Elias Zhang, using a two-stage LLM pipeline. First, gpt-5.1-2025-11-13 produced a chronologically consistent biographical framework spanning ages 15 to 64 (1960–2009) for a Chinese-American character, covering family background, geographic progression, education, career, family milestones, technology adoption, and cultural context. Second, conditioned on this framework and an evolving memory file summarizing previously generated years, gemini-2.5-pro-preview-05-06 generated 20 episodes per year (50 phases, 1,000 total) in first-person narrative style. The generation prompt targeted 50–60 words; the realized passages average 62.4 ± 4.8 words (95.1 ± 12.7 tokens). Each episode includes a specific temporal context, period-appropriate details, and a distinguishing theme to serve as a cue. Ten question–answer pairs were generated per episode with sufficient context to disambiguate among episodes. Nine paraphrases were additionally generated per episode. Full synthesis details are in Supplementary Note 1, and example episodes, questions and model outputs from every stage of the pipeline are collected in Supplementary Note 8.

#### Masked LoRA

We introduce a masked variant of LoRA (***Hu et al., 2022***) that restricts the low-rank update to a randomly selected subset of input and output neurons of a single weight matrix. Given a target weight matrix *W* ∈ ℝ*^d^*_out_^×*d*^_in_, standard LoRA reparameterizes the update as Δ*W* = *BA* with *B* ∈ ℝ*^d^*_out_^×*r*^, *A* ∈ ℝ*^r^*^×*d*^_in_. Our masked LoRA instead defines two index sets: *M*_in_ ⊂ {1, …, *d*_in_} and *M*_out_ ⊂ {1, …, *d*_out_}, each of cardinality *m*, selected uniformly at random and held fixed thereafter. The low-rank factors operate entirely in this *m*-dimensional subspace: *A* ∈ ℝ*^r^*^×*m*^, *B* ∈ ℝ*^m^*^×*r*^.

The forward pass for a masked LoRA adapter computes:

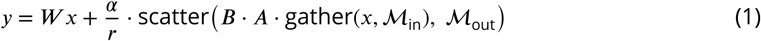

where gather(*x*, *M*_in_) ∈ ℝ*^m^* extracts the input coordinates indexed by *M*_in_, and scatter(⋅, *M*_out_) places the *m*-dimensional result back into the *d*_out_-dimensional output space (with zeros elsewhere). The total parameter count per adapter is 2*rm* = 4*m* (for *r* = 2). At the minimum mask sizes found in the capacity sweep, *m* scales approximately as half the passage length, corresponding to roughly 2 trainable parameters per stored token; the more conservative initialization used for the continual experiments corresponds to approximately 2.5 parameters per token.

The mask coordinates are sampled uniformly at random and then held fixed for the lifetime of the adapter, so the only trainable parameters are the *A* and *B* matrices. One further design choice is critical: the scaling factor 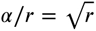 (with 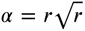) was found to stabilize training across varying mask sizes. We apply the adapter to the MLP down-projection at layer 10, identified as optimal by sweeping layers and target matrices (Fig. 4). Implementation details, including the masked forward pass and the mask auto-growth mechanism, are given in Supplementary Note 3.

### Adapter training and gating

For each new episode, a fresh masked LoRA adapter is initialized with a mask corresponding to approximately 2.5 parameters per token (initial *m* = 45 for Wiki and *m* = 60 for Character). The adapter is trained with AdamW (learning rate 0.05, no weight decay) for up to 1,000 epochs on the reconstruction objective: generate the stored passage given the common prompt ‘Reconstruct the story:’. Unlike the experiment in Fig. 1, the same reconstruction prompt is used for every episode. Training stops early upon achieving exact string match. If the initial mask is insufficient for perfect reconstruction, an auto-growth mechanism expands *m* by a factor of 1.2 and retrains (up to 10 retries).

Concurrently, each episode is embedded by Qwen3-Embedding-0.6B (***Zhang et al., 2025***; ***Yang et al., 2025***) (1024-dimensional, last-token pooling, L2-normalized) and stored alongside the adapter as a retrieval index. At inference, the incoming cue is embedded with the same encoder and the adapter whose stored passage embedding has the highest cosine similarity is selected. The index is part of the storage cost of the system: 1,000 episodes require 1.02 million embedding values (4.1 MB in float32). The realized float16 adapter stores occupy 0.39 MB for Wiki and 0.51 MB for Character, so the float32 index is approximately 10 and 8 times larger, respectively, though still only ∼0.013% of the base model’s parameter count. The encoder itself is shared across all episodes and is not a per-episode cost.

### Internal RAG

The iRAG pipeline operates in two stages. Stage 1 (routing and reconstruction): the incoming question is embedded, matched to the nearest stored episode embedding, and the corresponding adapter is composed with the base model to generate the stored passage (greedy decoding, max 150 new tokens). Stage 2 (QA): the adapter is removed, and the reconstructed passage is placed in the base model’s context followed by the question. The base model generates a short answer (greedy decoding, max 30 tokens) using standard in-context QA.

### Consolidation

#### Reconstruction-based consolidation

We update the base weights by fine-tuning with a high-rank LoRA (rank 2048, *α* = 4096) applied to MLP projections (down_proj, up_proj, gate_proj) across every transformer layer (32 for Llama and 36 for Qwen). This high-rank LoRA is used purely as a compute-efficient parameterization of a direct base-weight update. The learned increment is merged into the base matrices after training and then discarded. Training uses AdamW (learning rate 2.5 × 10^−5^, batch size 16, up to 21 epochs) distributed across 2 GPUs with FSDP. All 1,000 passages are trained simultaneously in batch mode. Consolidation, therefore, relearns the same episodes into the shared base weights by batch replay of the original passages, rather than transferring parameters from the episodic adapters. For embedding-cued consolidation, an unused reserved token in each model’s input template has its embedding replaced at the input layer with the passage’s Qwen3 embedding vector (1024-dim, zero-padded to 4096); for Llama this token is ID 128018. To enable question-cued recall without a stored index, Gaussian noise (per-element std sampled uniformly from [0, *σ*_max_], with *σ*_max_ = 0.039 for Wiki and 0.044 for Character) is added to passage embeddings during training, calibrated from the empirical 95th-percentile deviation between passage and question embeddings. Details in Supplementary Note 4.

#### Direct QA consolidation

Rather than training a consolidated model to reconstruct passages and then performing in-context QA, direct QA consolidation trains the model to answer questions about stored episodes directly from its parameters. We use the same high-rank base-weight update (LoRA rank 2048, *α* = 4096, every transformer layer, MLP targets, merged into the base weights after training) trained for 9 epochs with AdamW. We evaluate four training conditions: a passage-only baseline and three augmentation strategies of increasing aggressiveness (paraphrase, knowledge-instruction, and masked reconstruction). The latter two are each applied on top of paraphrase augmentation but are otherwise distinct recipes:

##### Condition A (passage only)

The base condition fine-tunes on passages in standard chat-template format, with the passage as the assistant’s response to a reconstruction prompt.

##### Condition B (+ paraphrases)

In addition to the original passage, 9 LLM-generated paraphrases are included as alternative reconstruction targets, diversifying the surface forms the model associates with each episode’s content.

##### Condition C (+ knowledge instruction)

Following the knowledge-instruction approach (***Ovadia et al., 2025***), declarative facts extracted from each passage are added as short question–answer pairs (prompts of the form “Tell me a fact about {title}” paired with an extracted sentence), up to 50 facts per episode rendered with varied templates (a value selected in a pilot grid search over the number of facts and of interleaved question–answer pairs). To teach the question-answering format without leaking test questions, auxiliary QA pairs drawn from held-out donor episodes and a name-anchoring drill are additionally included. This condition builds on paraphrase augmentation but does not include masked reconstruction.

##### Condition D (+ masked reconstruction)

A denoising objective is added in which 45% of tokens in the passage are replaced with an unused reserved token from the corresponding model tokenizer (ID 128013 for Llama), and the model is trained to reconstruct the original passage from the corrupted input. This masked autoregressive training forces the model to form robust internal representations rather than relying on surface-level pattern completion. Paraphrases are duplicated 3× in this condition to balance the additional masked examples. This condition is applied on top of paraphrase augmentation and does not include the knowledge-instruction facts. Implementation details are in Supplementary Note 5.

#### Validation/test split and epoch selection

Direct QA accuracy is not monotone in training epochs: with overfitting, the model can repeat the whole passage rather than answer the question naturally. The number of epochs must therefore be chosen per condition. We split the QA pairs of each episode into a validation set, used only to choose the number of training epochs, and a test set, used only to report accuracy. The validation/test split contains 3,179/2,694 unique question strings on Wiki and 2,868/2,343 on Character; because two retained Wiki question strings occur twice in the generated records, output-level analyses contain 2,696 Wiki test records per condition before excluding missing judge scores (Supplementary Note 6). For each condition we evaluate checkpoints at 1, 3, 6 and 9 training epochs, select the epoch with the highest validation accuracy, and report test accuracy at that epoch. Base-capability retention is measured on the consolidated model at that same epoch. At *N* = 1 the optimal epoch was chosen based on a pilot sweep of 20 episodes. All other hyperparameters (learning rate, rank, *α*, target modules) are fixed across conditions. Selected epochs and the corresponding accuracies are listed in Supplementary Note 5.

### Regularization baselines

Three regularization methods were evaluated on the continual fine-tuning setup at *T* = 100. L2: Penalizes the *L*_2_ distance of weights from their values at the start of each episode’s training. EWC (elastic weight consolidation) (***Kirkpatrick et al., 2017***): Penalizes changes to parameters weighted by the diagonal Fisher Information Matrix accumulated over all previous episodes. OPD (on-policy distillation) (***Li and Hoiem, 2018***): A KL divergence penalty between the model’s output distribution and a reference copy saved before each episode’s training, evaluated on 1,000 held-out Tulu 3 (***Lambert et al., 2024***) prompts (10 steps per episode, batch size 4, max 256 new tokens).

Regularization strength was swept over several orders of magnitude for each method.

### Evaluation

#### Reconstruction quality

An LLM judge (gpt-oss-120b (***OpenAI, 2025***), served via AWS Bedrock) evaluates free-form passage reconstructions. For the Character dataset, the judge extracts five fixed slots (event, social context, emotional takeaway, temporal context, spatial context) and scores each 0.0–1.0 for novel recall. For Wikipedia, the judge dynamically identifies 4–8 informational aspects and scores each 0.0–1.0. The reported “recall judge score” is the mean across slots/aspects, adjusted for content guessable from the title alone. The judge prompts and analysis on the evaluation metrics are in Supplementary Note 6; Supplementary Fig. 4 compares the resulting forgetting curves with surface-based metrics.

#### QA accuracy

A binary LLM judge (the same gpt-oss-120b model) evaluates short-form answers against reference answers. Answers are scored as correct if they match the reference in meaning, even if paraphrased or containing additional context; they are scored as incorrect only if the core fact is fundamentally wrong or the answer is empty/refusal. An answer was counted as correct only if the judge marked it correct *and* the response contained at most one sentence; paraphrasing and additional context within that sentence were permitted. Sentences are counted by splitting on sentence-final punctuation followed by whitespace. The length restriction matters because a consolidated model often answers by reconstructing the whole episode, which may contain the reference fact incidentally. QA pairs are generated from the episode passages with gpt-5.2-2025-12-11. We then remove two classes of question: (i) those the base model already answers correctly without any fine-tuning, for which a correct answer is not evidence of storage or retrieval, and (ii) those that are not answered correctly even when the source passage is placed verbatim in the context window (the *N* = 1 in-context condition), which are defective items rather than hard memory problems. The filter is defined by this in-context reference condition and is applied identically to every method and every *N*. Both criteria are evaluated for each of the two base models in turn, and a question is kept only if it survives all four; this retains 5,211 Character and 5,873 Wiki questions, with pre-existing knowledge accounting for most of the removals. Supplementary Note 2 reports the counts by stage and accuracy on the complete, unfiltered question set; no conclusion in this paper depends on the filter. Supplementary Note 6 compares this judge against token F1, ROUGE-1, and ROUGE-L on the answers behind Fig. 9C. The ranking of the consolidation conditions is unchanged under every one of them.

#### Uncertainty

Measurement uncertainty on the question-answering figures is quantified by a cluster bootstrap over episodes: episodes are resampled with replacement 2,000 times, all questions belonging to a resampled episode are taken together, and we report the 2.5th and 97.5th percentiles of the resulting distribution. Clustering by episode rather than by question is the conservative choice, since questions from the same episode share a passage, an adapter and a routing decision and are therefore not independent. Because iRAG accuracy is the *N* = 1 accuracy propagated through top-1 routing, and the former is 100% by construction, the interval on iRAG accuracy coincides with the interval on routing accuracy. Experiments at *N* = 1,000 use the complete dataset and one training run unless a caption states otherwise; the bootstrap intervals therefore quantify evaluation-sampling uncertainty, not between-training-seed variability. Figure captions specify the subset repetitions used at smaller *N*.

#### General capability retention

We track WinoGrande (***Sakaguchi et al., 2021***), HellaSwag (***Zellers et al., 2019***), and MMLU (***Hendrycks et al., 2021***) (zero-shot, no chat template) throughout training, normalized by the base model’s scores.

## Supplementary information

Supplementary File 1 accompanies this article. It contains Supplementary Figures 1–12 and Supplementary Notes 1–8 (Character dataset synthesis, question filtering, masked LoRA implementation, reconstruction-based consolidation, direct QA consolidation implementation, LLM judges and evaluation metrics, results on Qwen3-8B, and examples).

## Acknowledgments

This work was supported by the Kempner Institute for the Study of Natural and Artificial Intelligence at Harvard University, the Office of Naval Research (ONR) grant No. N00014-23-1-2051, the Gatsby Charitable Foundation, and an Amazon Research Award Spring 2025. Any opinions, findings, and conclusions or recommendations expressed in this material are those of the authors and do not reflect the views of Amazon. We have benefited from very helpful discussions with Aya Ben Yakov, Jorin Overwiening, Qianyi Li, Jingxuan Fan, and Isaiah Kletenik.

## Author contributions

X.P. designed the framework, implemented the code, performed the analysis, and wrote the manuscript. E.H. designed the dataset and conducted the experiments. R.S. contributed to the consolidation experiments and replicated the Llama experiments on Qwen. H.S. supervised and managed the project and provided the key design ideas.

## Data availability

The two datasets used in this study, along with their generation code, are available at https://github.com/xup5/Continual-Episodic-Memory-in-Large-Language-Models.

## Code availability

All code required to reproduce the experiments and figures, including the masked LoRA implementation, the gating and internal-RAG pipelines, the consolidation training scripts, and the evaluation and plotting code, is available at https://github.com/xup5/Continual-Episodic-Memory-in-Large-Langu age-Models.

## Supplementary Information

## Supplementary Figures

**Figure 1:**
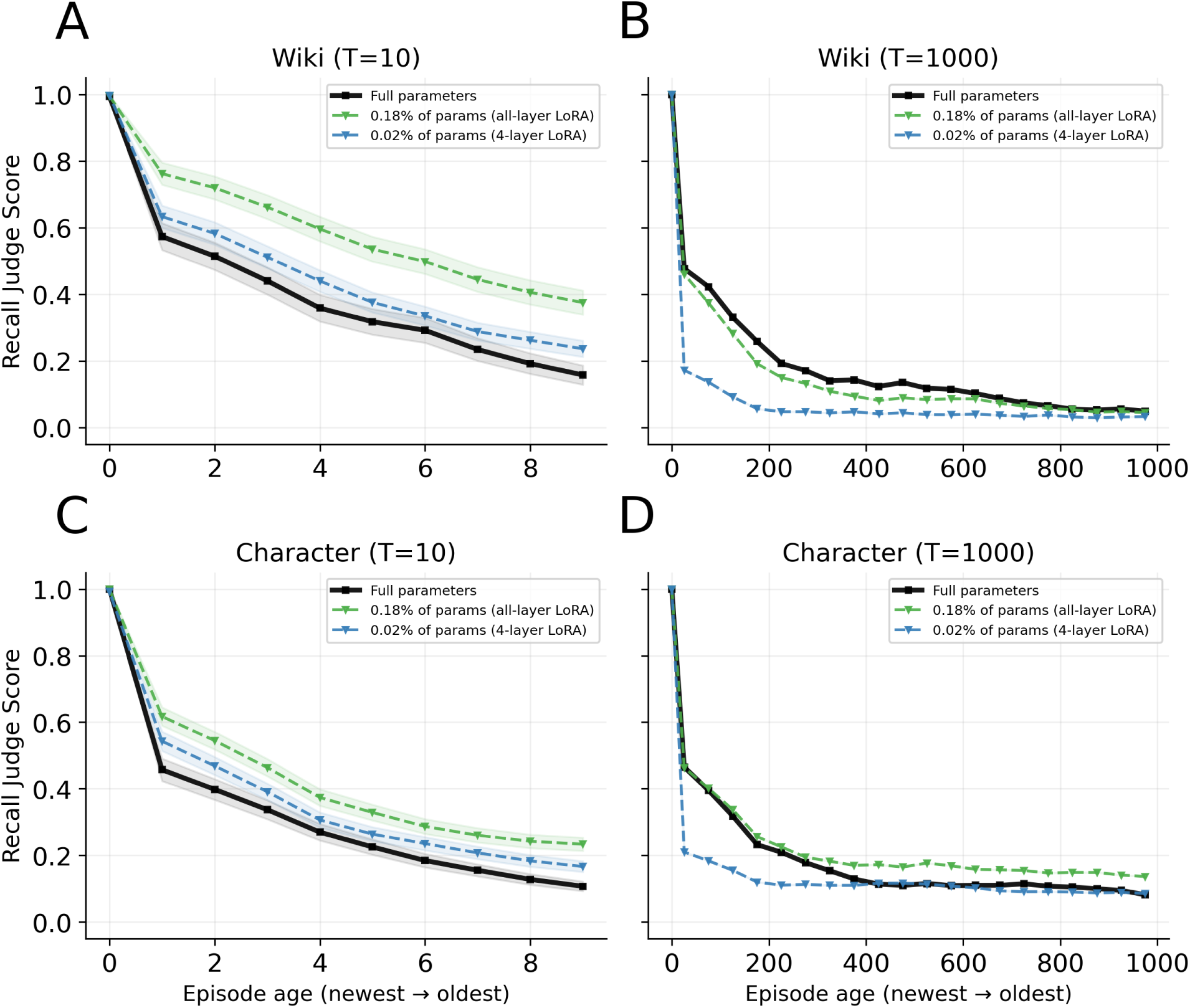
Continual-learning forgetting curves for streams of *T* = 10 and *T* = 1,000 sequential episodes, complementing the *T* = 100 results in main-text Fig. 1. The *T* = 1,000 panels reproduce the corresponding main-text panels to permit side-by-side comparison with *T* = 10. The same qualitative pattern holds at both extremes: full-parameter fine-tuning (solid) forgets fastest, while LoRA variants using 0.176% (all-layer, green dashed) and 0.022% (4-layer, blue dashed) of the parameters slow forgetting but do not prevent it.

**Figure 2:**
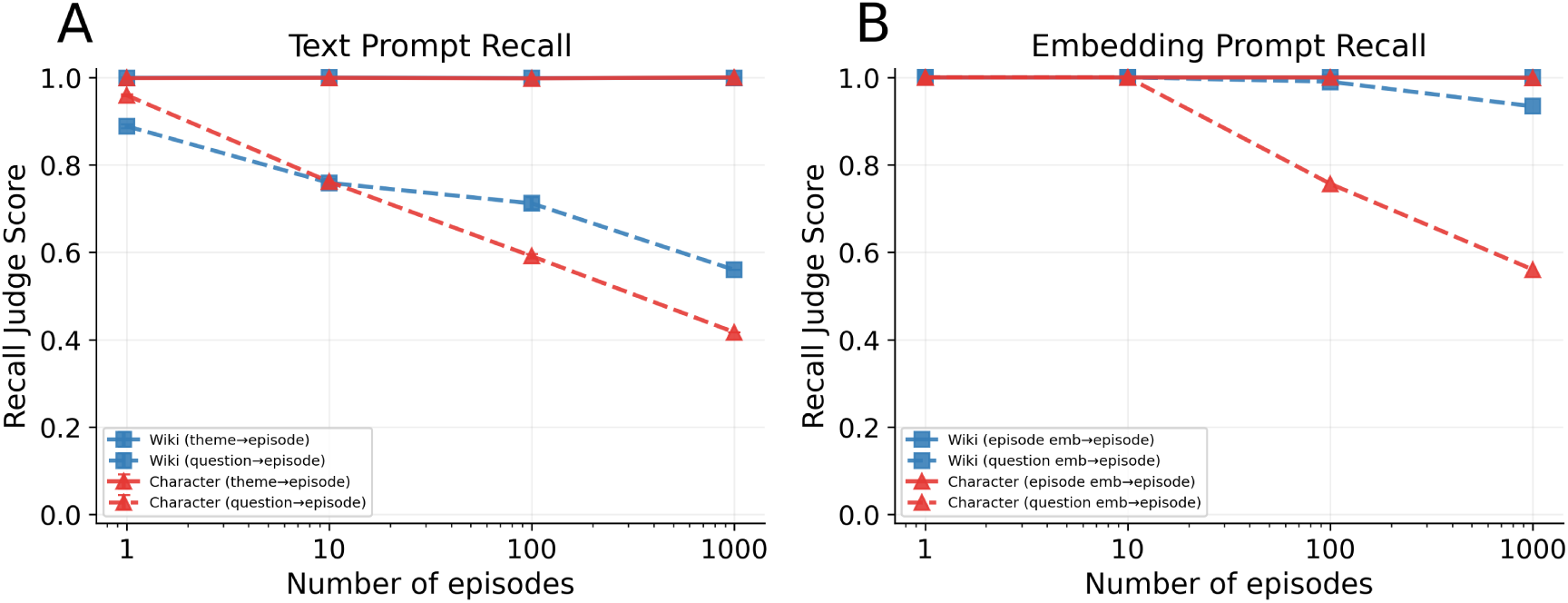
Reconstruction-based consolidation. **(A)** Text-prompted recall: the consolidated model reconstructs the full episode passage from a text cue. Theme-cued recall (solid) remains near-perfect across all library sizes including *N* =1,000. Question-cued recall (dashed) degrades with scale, as the question prompt provides only a partial cue and the model must disambiguate among overlapping episodes. **(B)** Direct embedding-cued recall: the passage embedding (solid) or the question embedding itself (dashed) is injected as input, with no nearest-episode routing step. Passage-embedding recall remains near-perfect, demonstrating that a single dense vector carries sufficient information for faithful reconstruction. Direct question-embedding cue recall, enabled by noise-augmented training, degrades at scale on the Character corpus. This mechanism is distinct from the routed question-embedding cue in main-text Fig. 8C.

**Figure 3:**
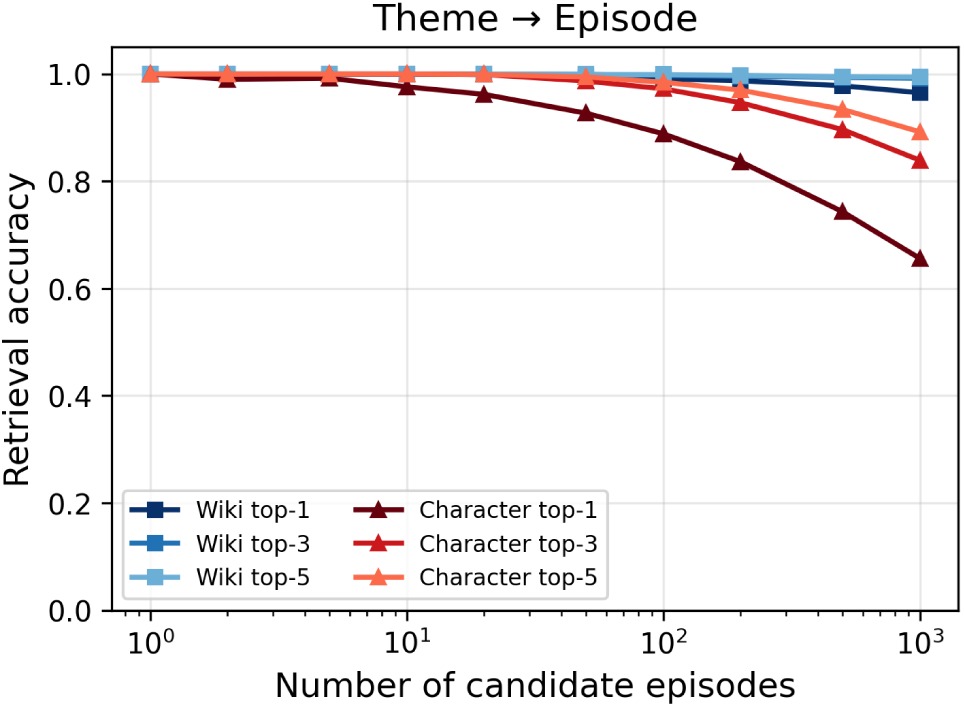
Theme-to-episode retrieval accuracy (top-1, top-3, top-5) as a function of the number of candidate episodes. The embedding of the theme is used to select the gated LoRA adapters based on the match to the episode embeddings. Theme cues carry less semantic information than questions, leading to substantially lower routing accuracy on the Character corpus. At *N* =1,000, Character top-1 accuracy is 65.6%, compared to 96.5% for Wiki. Top-5 retrieval partially recovers accuracy (89.2% Character, 99.4% Wiki).

## Supplementary Note 1 – Character Dataset Synthesis

The synthetic character corpus centers on a fictional character, Elias Zhang, generated using a two-stage LLM pipeline designed to produce a temporally coherent, semantically overlapping set of autobiographical episodes.

### Stage 1: Biographical Framework

A biographical framework was generated using gpt-5.1-2025-11-13. The character is a Chinese-American born in 1945 in San Francisco. The framework covers ages 15 to 64, spanning the years 1960–2009, and provides a year-by-year chronological outline including: family background, geographic progression, education, career milestones, family events, technology adoption, and cultural touchstones. Each life stage is grounded in the historical and cultural context of the corresponding era, covering diverse life domains (work, travel, relationships, hobbies, health).

### Stage 2: Episode Generation

Episodes were generated using gemini-2.5-pro-preview-05-06 via the Harvard Gemini API (temperature = 0.7, topK = 40, topP = 0.95). The corpus comprises 1,000 stories across the 50 years of the chronology (20 stories per year), written in first-person narrative style. Generation was conditioned on both the biographical framework and an evolving memory file that summarizes previously generated years, ensuring cross-year consistency.

The generation prompt targeted 50–60 words per story; the realized corpus averages 62.4 *±* 4.8 words (mean *±* s.d.; 95.1 *±* 12.7 Llama tokens). Each story is written in a voice appropriate to Elias’s age and constrained to include specific temporal context, period-appropriate details, and a distinguishing theme suitable for use as a retrieval cue. Ten question-answer pairs were generated per story in a later pass with gpt-5.2-2025-12-11 (factual, second-person phrasing, answers of at most five words, with sufficient context in the question to disambiguate among episodes).

Additionally, 9 paraphrases were generated per story using gpt-5.2-2025-12-11. Four generation phases (14, 28, 34, 39) failed on the first attempt and were regenerated.

## Supplementary Note 2 – Question Filtering

Both benchmarks were generated automatically, and QA accuracy is scored by an LLM judge. Two failure modes therefore contaminate raw accuracy in ways that have nothing to do with whether a memory system stored the episode. First, a question may be answerable from the base model’s pre-training knowledge alone, in which case a correct answer is not evidence of successful storage or retrieval. Second, a question may be unanswerable from its own source passage — the generator produced an ambiguous or under-specified question, the reference answer is wrong, or the judge cannot score it reliably — in which case an incorrect answer is not evidence of failed storage or retrieval.

We identify the second class operationally. With a single stored episode (*N* = 1) there is no retrieval problem to solve: the correct passage is placed verbatim in the context window, and the model answers by ordinary in-context reading. This condition is our *gold standard* : the ceiling that any memory system can aspire to, and the same reference point used when retrieval-augmented generation is evaluated with an oracle retriever. A question that the model cannot answer even when handed its own source passage is a defective item, not a hard memory problem. We therefore exclude it, together with the “stale” items, and report accuracy on the remaining questions.

Two consequences of this design should be stated plainly. (i) The exclusion is defined by a condition of the evaluation protocol (*N* = 1 in-context reading), not by the performance of any system we propose; the same filter is applied identically to direct QA, iRAG, and all consolidation conditions, and to both the adapter and the consolidated-weight pathways. (ii) Because the filter retains only questions answered correctly from their source passage, iRAG accuracy at *N* = 1 is 100% on the retained set. The informative quantity is its degradation as the number of candidate episodes increases and routing becomes nontrivial.

### Applying both criteria to both base models

Both criteria depend on a base model, and this study uses two. They are therefore applied twice, in sequence. Starting from the 10,000 generated questions per corpus, evaluating with Llama leaves 6,669 (Wiki) and 6,350 (Character); evaluating again with Qwen3-8B removes a further 668 and 888 questions that base Qwen already answers, and 128 and 251 that Qwen cannot answer even with the source passage in context. The final sets contain 5,873 Wiki and 5,211 Character questions. Unless explicitly labeled as an unfiltered sensitivity analysis in this note, all QA results in the main text and supplementary figures use these final filtered sets.

A retained question is therefore one that neither base model can answer from pre-training alone and that both can answer when handed its source passage; it isolates memory rather than prior knowledge or benchmark defects. Because the *N* = 1 criterion is applied to both models, the identity noted in (ii) above holds for both: iRAG accuracy at *N* = 1 is 100% by construction in Fig. 7C and in Supplementary Fig. 10C, and the informative quantity in each is the degradation to *N* = 1,000.

The two models agree only partly about what counts as already known: of the 3,923 Wiki questions stale for at least one model, 2,163 are stale for both, and the corresponding Character figures are 4,153 and 1,746. These union counts also include 13 Wiki and 43 Character questions that Qwen answers from pre-training but that had already been removed as Llama *N* = 1 failures; they therefore do not appear in the sequential “Stale (Qwen3-8B)” row of Table 1. The partial agreement is expected from different pre-training corpora and cutoffs, and it is why filtering on one model alone left a residue of questions the other could already answer. Removing that residue brings the two direct-QA floors into the same range (Wiki 1.3% for Llama against 4.4% for Qwen, where filtering on Llama alone left 1.8% against 8.4%); the remaining difference reflects genuine differences in zero-shot guessing rather than an artifact of the filter.

### Number of excluded questions by category

Each corpus contains 1,000 episodes with 10 questions each, i.e. 10,000 questions (Wiki contains 18 duplicated question strings, leaving 9,982 unique items). Table 1 reports the counts stage by stage. Pre-existing base-model knowledge accounts for most of the exclusions: 32.5% of Wiki and 32.2% of Character questions are already answered by Llama, and a further 668 and 888 are newly removed as stale under Qwen3-8B among the Llama survivors. Before overlap with the stale set is removed, Llama flags 102 Wiki and 489 Character questions as *N* = 1 failures; 71 and 428, respectively, are newly removed and therefore appear in the table. Qwen is then evaluated only on the Llama survivors and newly removes 128 Wiki and 251 Character *N* = 1 failures. The higher Character rate for this criterion reflects the narrative form of that corpus, in which some generated questions presuppose surrounding context that the individual passage does not in fact contain. In total the filter retains 5,873 of 9,982 Wiki (58.8%) and 5,211 of 10,000 Character questions (52.1%).

**Table 1:** Questions excluded from QA accuracy computation. The filter is applied sequentially: the two criteria are first evaluated on Llama, then re-evaluated on Qwen3-8B, and the surviving set is used for every QA result except the explicitly labeled sensitivity analysis in Table 2. “Stale” = answered correctly by that base model with no fine-tuning; “*N* = 1 fail” = not answered correctly with the source passage in context. Counts are unique questions; Wiki has 9,982 unique strings among its 10,000 pairs.

| Stage | Character |  | Wiki |  |
| --- | --- | --- | --- | --- |
|  | removed | remaining | removed | remaining |
| All generated questions | — | 10,000 | — | 9,982 |
| Stale (Llama) | 3,222 |  | 3,242 |  |
| $N=1$ fail (Llama)<sup>†</sup> | 428 | | 71 | |
| after Llama filter |  | 6,350 |  | 6,669 |
| Stale (Qwen3-8B) | 888 | 5,462 | 668 | 6,001 |
| $N=1$ fail (Qwen3-8B) | 251 | 5,211 | 128 | 5,873 |
| **Retained for scoring** |  | **5,211** (52.1%) |  | **5,873** (58.8%) |

<sup>†</sup> Questions already removed as stale are not counted again, so each column sums to the remaining total. The raw numbers of $N=1$ failures, before removing that overlap, are 489 (Character) and 102 (Wiki).

### Sensitivity analysis: accuracy on the complete question set

Table 2 reports the principal QA numbers under three scoring sets: the complete question set, the set with only stale questions removed, and the filtered set used in the main text. No conclusion in the paper depends on the filter.

**Table 2:** QA accuracy under three scoring sets, all for Llama. “Complete” uses all questions; “stale removed” excludes only questions Llama already answers; “filtered” is the set used in the main text. Sample sizes and values are given as Character/Wiki.

| Dataset | Condition | Complete<br>($n=10,000/9,982$) | Stale removed<br>($n=6,778/6,740$) | Filtered (main text)<br>($n=5,211/5,873$) |
| --- | --- | --- | --- | --- |
| Character | iRAG, $N=1$ | 95.1% | 93.7% | 100% |
| | iRAG, $N=1,000$ | 81.1% | 79.9% | 85.4% |
| | Direct QA, $N=1,000$ | 2.8% | 1.0% | 0.8% |
| Wiki | iRAG, $N=1$ | 99.0% | 98.9% | 100% |
| | iRAG, $N=1,000$ | 98.4% | 98.4% | 99.3% |
| | Direct QA, $N=1,000$ | 6.5% | 1.8% | 1.3% |

On the complete question set (*n* = 10,000 Character and 9,982 Wiki), iRAG reaches 95.1% and 99.0%, respectively, at *N* = 1, and 81.1% and 98.4% at *N* = 1,000. On the filtered set used in the main text, the corresponding values are 100% at *N* = 1 and 85.4%/99.3% at *N* = 1,000. Relative to the complete-set analysis, filtering changes the reported *N* = 1,000 values by approximately 4.3 and 0.9 percentage points. Direct QA moves in the opposite direction: complete-set accuracy is higher (2.8% Character and 6.5% Wiki at *N* = 1,000) than filtered accuracy (0.8% and 1.3%), because the stale questions removed by the filter are precisely those the base model could already answer. Removing them makes the direct-QA failure look worse, not better. Thus, the large gap between direct QA and iRAG holds under every scoring set.

The same filter is used for both base models (Supplementary Note 7), so these unfiltered accuracies are the relevant sensitivity analysis for both: the effects reported in the paper survive when no filter is applied at all.

## Supplementary Note 3 – Masked LoRA Implementation

### Mask construction

For a target weight matrix *W ∈* R*^d^*_out_*^×d^*_in_ (in our case the MLP down-projection, *d*_out_ = 4096, *d*_in_ = 14336), masked LoRA defines two index vectors *M*_in_ *∈* {0*, …, d*_in_ *−* 1}*^m^* and *M*_out_ *∈* {0*, …, d*_out_ *−* 1}*^m^* by sampling *m* coordinates uniformly at random (without replacement) from each dimension.

### Forward pass

Given input *x ∈* R*^d^*^in^, the adapter forward pass is:

1. **Gather:** *x*_sub_ = index select(*x, M*_in_) *∈* R*^m^*
2. **Down-project:** *h* = *A · x*_sub_ *∈* R*^r^*, where *A ∈* R*^r×m^*
3. **Up-project:** *y*_sub_ = *B · h ∈* R*^m^*, where *B ∈* R*^m×r^*
4. **Scatter:** index add (**0***_d_*_out_ *, M*_out_*, y*_sub_ *· s*)

where the scaling factor *s* = *α/r* with 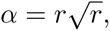, giving 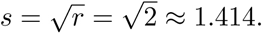

### Initialization

Matrix *A* is initialized with Kaiming uniform (fan-in = *m*), and *B* is initialized to zeros, so that the adapter output is zero at initialization and training begins from the pretrained model’s behavior. This follows the standard LoRA initialization convention.

### Auto-growth mechanism

The initial mask size is set to approximately 2.5 trainable parameters per passage token (*m* = 45 for Wikipedia passages averaging 76 tokens and *m* = 60 for Character passages averaging 95 tokens; an adapter contains 4*m* parameters). If training does not achieve exact string match of the target passage within the maximum epoch budget, the mask size is grown by a factor of 1.2 and training restarts from fresh initialization, up to 10 retries.

### Parameter efficiency

With *r* = 2, the realized 1,000-episode stores contain approximately 197,000 adapter parameters for Wiki and 256,000 for Character. Stored in float16, these occupy approximately 0.39 MB and 0.51 MB, respectively, negligible compared with the 16 GB base model.

The complete episodic store has a second component. Each episode is also indexed by a 1,024-dimensional embedding used for routing, so the index holds 1,000 *×* 1,024 = 1.02 million values, or 4.1 MB as stored here in float32 (2.0 MB at half precision). The float32 index is therefore approximately 10 times larger than the Wiki adapters and 8 times larger than the Character adapters, and it, rather than the adapters, dominates the per-episode storage. Both components remain small against the base model: the adapters are *∼*0.003% of its parameter count and the index *∼*0.013%. The “two to three parameters per token” figure quoted in the main text refers to trainable adapter parameters; the index is not trained, and the shared sentence encoder that produces it (Qwen3-Embedding-0.6B) is a fixed component of the retrieval mechanism rather than per-episode storage.

## Supplementary Note 4 – Reconstruction-Based Consolidation

Reconstruction-based consolidation transfers the ability to reconstruct stored passages from per-episode adapters into the base model’s own parameters.

### Training setup

The base model is fine-tuned with LoRA (rank 2048, alpha 4096) applied to MLP projections (down proj, up proj, gate proj) across every transformer layer (32 for Llama and 36 for Qwen). Training uses AdamW with learning rate 2.5 *×* 10*^−^*^5^, batch size 16, for up to 21 epochs. Training is distributed across 2 GPUs using FSDP (Fully Sharded Data Parallelism).

### Embedding injection mechanism

An unused reserved token from the corresponding model tokenizer is placed in the input template (for Llama, token ID 128018, <|reserved special token 10|>). During forward passes, the token’s embedding in the input embedding layer is replaced with the passage’s Qwen3-Embedding-0.6B vector (1024 dimensions, zero-padded to 4096 to match the model’s hidden size). This allows the model to condition reconstruction on a dense semantic cue rather than a text prompt.

### Noise-augmented training

To enable question-embedding-cued reconstruction (where the model receives a question embedding at inference but was trained on passage embeddings), Gaussian noise is added to passage embeddings during training. The per-element standard deviation is sampled uniformly from [0*, σ*_max_] where *σ*_max_ = 0.0389 (Wikipedia) or 0.0435 (Character). These values were calibrated from the empirical 95th-percentile deviation between passage and question embeddings in the training corpus.

## Supplementary Note 5 – Direct QA Consolidation Implementation

This section provides implementation details for the four direct QA consolidation conditions described in the Methods.

### General setup

All four conditions share the same optimization: a rank-2048 LoRA (*α* = 4096) on the three MLP projections of every transformer layer, trained with AdamW at learning rate 2.5 *×* 10*^−^*^5^ and batch size 16 for 9 epochs. The conditions differ only in what the model is trained to produce, described below. Each condition is evaluated at two library sizes: a single episode per model, so that 1,000 separate models are trained and their accuracies averaged, and all 1,000 episodes consolidated into one model.

### Scoring

QA accuracy is computed on the filtered question set (Supplementary Note 2), and an answer counts as correct only if the judge marks it correct and it is at most one sentence long. The length restriction matters because the consolidated model often answers by reconstructing the whole episode rather than stating the fact. Such an answer may contain the reference fact incidentally, and crediting it would measure reconstruction rather than the direct question answering at issue here; Supplementary Note 8 gives examples.

We split each episode’s questions in half: a validation half used only to choose the optimal training epochs, and a test half used only for reporting accuracy: 3,179 validation and 2,694 test questions on Wiki, and 2,868 and 2,343 on Character. For each condition we evaluate the checkpoints at 1, 3, 6 and 9 epochs, take the epoch with the highest validation accuracy, and report accuracy on the test half at that epoch.

At *N* = 1 the same principle applies but the sweep runs on a subset of episodes. The epoch was chosen on a separate sweep of 20 evenly spaced episodes (0, 50, …, 950), and we report test-half accuracy over the remaining episodes.

**Table 3:** Epochs selected on the validation half, and the test-half accuracies reported at those epochs. (*N* = 1,000; main-text Fig. 9C and Supplementary Fig. 12C). Brackets give cluster-bootstrap 95% confidence intervals over episodes (2,000 resamples). Selection used only validation questions; the accuracies shown use only test questions.

| Condition | Wiki |  | Character |  |
| --- | --- | --- | --- | --- |
|  | Epoch | Test accuracy (%) | Epoch | Test accuracy (%) |
| *Llama-3.1-8B-Instruct* |  |  |  |  |
| No augmentation | 6 | 31.0 [29.3, 32.8] | 3 | 32.9 [31.0, 35.0] |
| Paraphrase | 1 | 44.5 [42.7, 46.5] | 1 | 40.2 [38.0, 42.2] |
| Knowledge instruction | 6 | 73.1 [71.4, 74.9] | 3 | 50.5 [48.5, 52.5] |
| Masked reconstruction | 3 | 69.1 [67.2, 71.0] | 9 | 59.2 [57.3, 61.3] |
| *Qwen3-8B* |  |  |  |  |
| No augmentation | 9 | 18.7 [17.1, 20.2] | 6 | 23.3 [21.6, 25.1] |
| Paraphrase | 3 | 48.1 [46.1, 49.9] | 3 | 40.8 [38.8, 42.9] |
| Knowledge instruction | 1 | 58.1 [56.3, 60.1] | 3 | 38.5 [36.6, 40.5] |
| Masked reconstruction | 9 | 59.5 [57.5, 61.4] | 6 | 44.9 [42.9, 46.8] |

### Condition A: Passage only

Each of the 1,000 episodes is formatted as a single chat-template exchange:

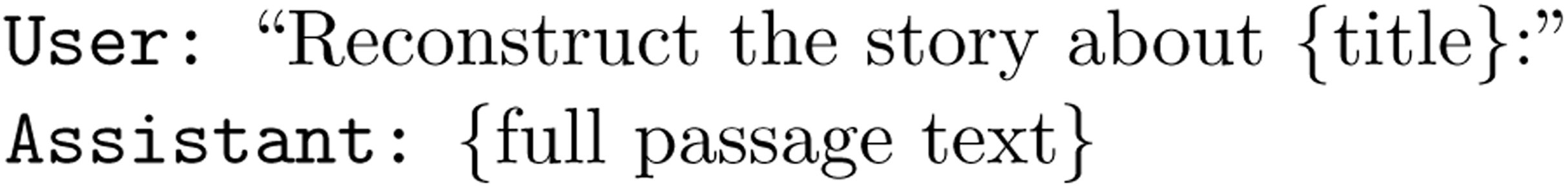

This establishes the baseline: whether LoRA fine-tuning alone can internalize passage content for downstream QA without any explicit QA supervision.

### Condition B: Paraphrase augmentation

In addition to the original passage, 9 LLM-generated paraphrases (produced by gpt-5.2-2025-12-11) serve as alternative assistant responses to the same reconstruction prompt. The training set thus contains 10 variants per episode. Paraphrases emphasize word-level substitution, sentence-level reordering, and paragraph-level restructuring while preserving all factual content. This diversification encourages the model to encode semantic content rather than memorizing specific surface forms.

### Condition C: Knowledge instruction

Following the knowledge-instruction paradigm [1], this condition supplements paraphrase reconstruction with QA-format supervision built from declarative facts. For each episode, the passage and its paraphrases are split into sentences of at least five words; these are deduplicated and capped at 50 facts per episode (on average *∼*25 facts per episode). Each fact is rendered as a chat exchange in which the user prompt is one of 25 templates of the form “Tell me a fact about *{*title*}*” and the assistant response is the extracted sentence; each fact is instantiated with 3 randomly chosen templates. The training mixture additionally includes (i) 30 auxiliary QA pairs drawn from held-out donor episodes (detailed below), which teach the short-answer QA format without exposing the memorized episodes’ own test questions, and (ii) a name-anchoring drill that maps an episode description back to its title. The system prompt is a generic assistant prompt for Wiki and an in-character persona for the Character dataset; cross-entropy loss is masked to the assistant turn. Unlike Condition D, this condition does not apply masked reconstruction.

To keep the auxiliary QA disjoint from the memorized episodes, the auxiliary QA pairs are drawn from a fixed pool of 20 donor episodes held out entirely from the consolidated set (i.e. outside the *N* episodes being memorized), rather than from the memorized episodes themselves. Because the donors are external, the *N* = 1,000 run consolidates the full 1,000 episodes, exactly matching Conditions A, B, and D, while remaining leak-free by construction. An earlier within-batch scheme, in which auxiliary QA was instead sampled from the memorized episodes, leaked test questions at *N* = 1,000 and inflated accuracy to an implausible *∼*98%; all reported knowledge-instruction numbers use the external-donor protocol.

### Condition D: Masked autoregressive reconstruction

A denoising pre-training objective is added. For each passage (and its paraphrases), a corrupted version is constructed by independently replacing each token with probability *t* = 0.45 with an unused reserved token from the corresponding model tokenizer (for Llama, ID 128013, <|reserved special token 5|>).

The mask is applied only to passage tokens in the user turn; instruction tokens are preserved. The training example is formatted as:

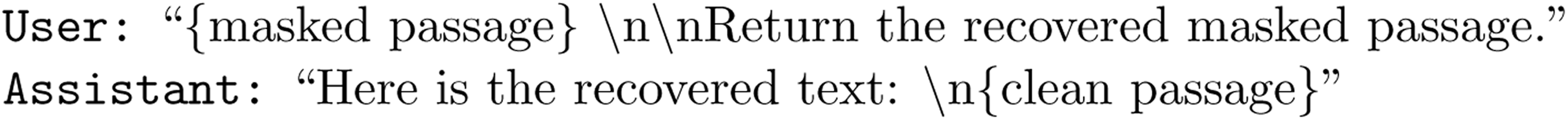

The masking probability *t* = 0.45 was chosen as a fixed value (not scheduled or sampled) to provide substantial corruption while retaining enough context for reconstruction. In this condition, paraphrases are duplicated 3*×* to balance the training distribution against the additional masked examples. A fresh mask is sampled independently for each training example at each epoch.

This denoising objective forces the model to develop robust internal representations of passage content: rather than relying on autoregressive continuation from the preceding tokens, the model must reconstruct masked spans from the global context of the passage, encouraging deeper semantic encoding.

## Supplementary Note 6 – LLM Judges and Evaluation Metrics

Two distinct LLM-based judges are used for evaluation.

### Reconstruction judge (recall evaluation)

An LLM judge (OpenAI gpt-oss-120b, served via AWS Bedrock) evaluates the faithfulness of free-form passage reconstruction. For the character dataset, the judge extracts five fixed slots (event, social context, emotional takeaway, temporal context, spatial context) and scores each on a scale of 0.0–1.0 for novel recall (content not guessable from the title alone). For the Wikipedia dataset, the judge dynamically identifies 4–8 informational aspects and scores each 0.0–1.0. The final score is the mean across all slots or aspects.

### QA judge

The same model scores short-form question answering, as a binary judge. The Methods summarize the criterion; the prompt is reproduced here verbatim, since the criterion is defined by it.

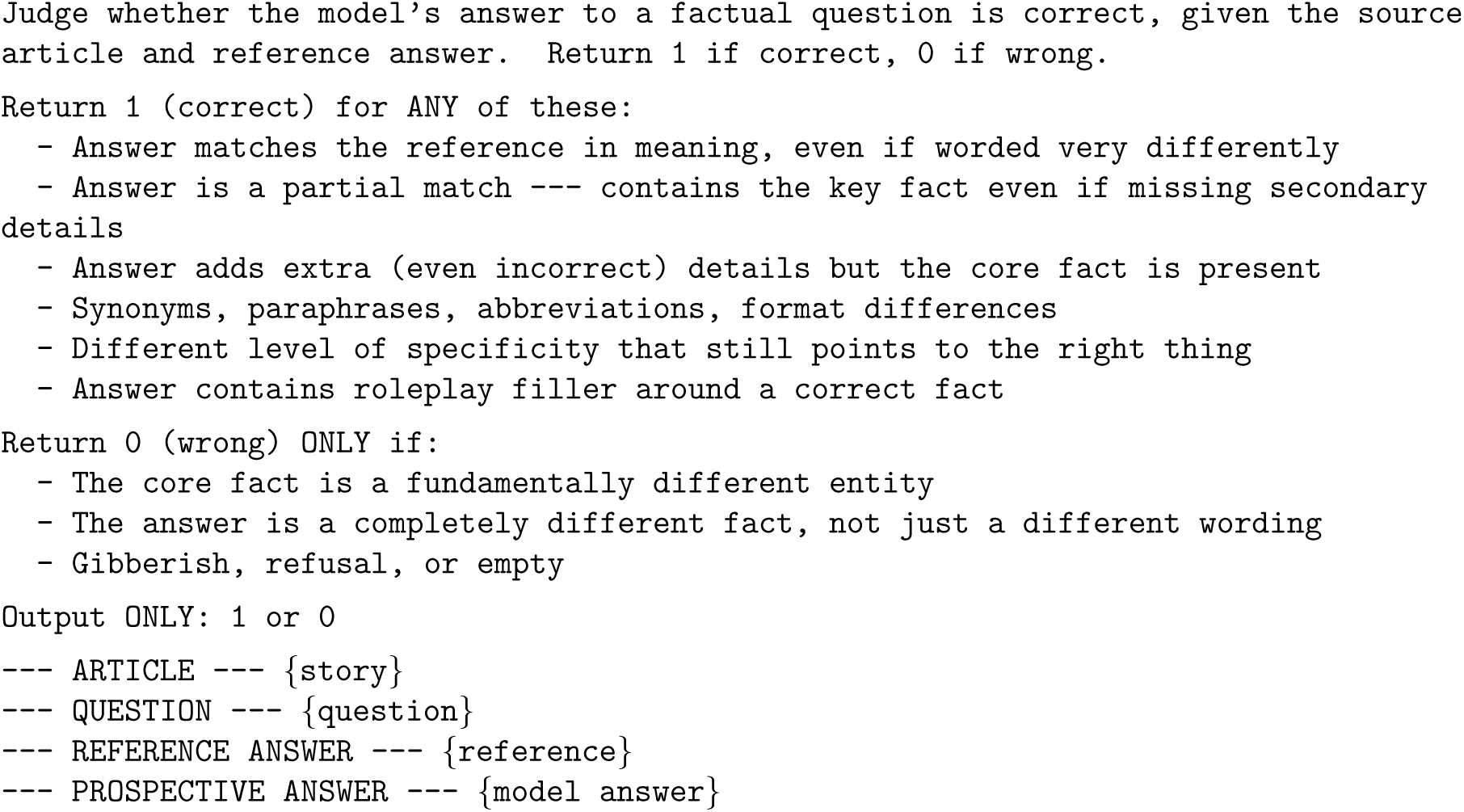

The criterion credits the reference fact alone, so a model is not penalized for paraphrasing or for supplying context around a correct answer.

String matching is not an adequate substitute for this judge. Table 4 compares the two scoring methods on the same generations: they agree to within 0.1 pp for plain full fine-tuning, whose answers echo the reference wording, but string matching understates accuracy by 30–38 pp for LoRA-tuned and system-prompted variants, whose answers paraphrase. Since the conditions compared in this paper differ exactly along that axis, string-match scores would not be comparable across them.

**Table 4:** Why QA is scored by an LLM judge rather than by string matching. Comparison on the Character corpus at *N* =50 episodes, 100 random subsamples per method, using batch fine-tuning. Rough match agrees with the LLM judge for plain full fine-tuning, but underestimates accuracy by 30–38 absolute points for LoRA-tuned and system-prompted variants, whose answers paraphrase the ground truth. “Gap” is the LLM-judged minus rough-match accuracy.

| Method | Rough match (%) | LLM-judged (%) | Gap (pp) |
| --- | --- | --- | --- |
| Full fine-tuning | 54.3 | 54.2 | −0.1 |
| Full fine-tuning + system prompt | 26.2 | 56.6 | +30.4 |
| LoRA-MLP ($r=2048$, $\alpha=4096$) | 31.9 | 65.7 | +33.8 |
| LoRA-MLP + system prompt | 28.9 | 66.9 | +38.0 |

### Comparison with surface metrics

We evaluate free-form passage reconstruction with four metrics: teacher-forced token accuracy (TF Acc), Levenshtein ratio, ROUGE-L, and an LLM judge. TF Acc measures the fraction of tokens the model predicts correctly when the ground-truth prefix is provided at each step; it reflects how well the model has memorized the passage during training but does not test whether it can produce the passage autoregressively. Levenshtein ratio and ROUGE-L measure character-level edit-distance similarity and longest-common-subsequence word overlap, respectively; both are purely surface-level. The LLM judge decomposes each passage into informational aspects (for Wikipedia) or narrative slots (for Character) and scores each on a 0–1 scale for the content that could not be guessed from the title alone.

Figure 4 compares all four metrics on the continual-learning forgetting curves. Two patterns stand out. First, TF Acc remains high (*>* 0.8) long after generation-based metrics have collapsed, confirming that teacher forcing masks catastrophic forgetting: the model can predict the next token given the correct prefix but cannot reconstruct the passage autoregressively. Second, Levenshtein and ROUGE-L track each other closely but diverge from the LLM judge, particularly on the Character corpus, where shared vocabulary across episodes (the recurring character Elias Zhang, recurring locations, recurring social relationships) inflates surface overlap even when the semantic content is wrong.

**Figure 4:**
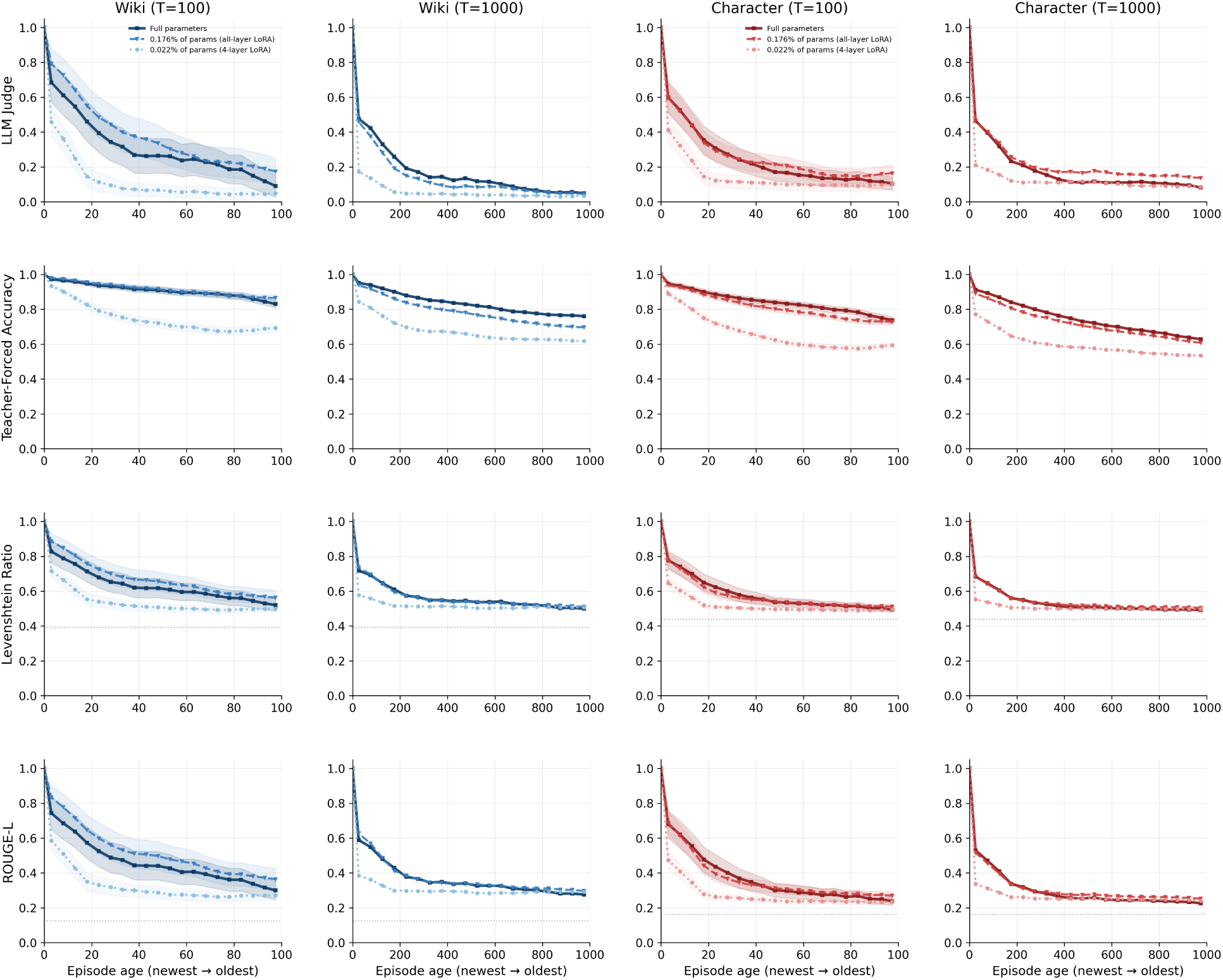
Forgetting curves measured by four reconstruction metrics across two datasets (Wikipedia, Character) and two sequence lengths (*N* =100, *N* =1,000). Rows from top to bottom: LLM judge, teacher-forced token accuracy (TF Acc), Levenshtein ratio, ROUGE-L. Each panel shows three architectures: full-parameter fine-tuning (solid), all-layer LoRA (dashed), and 4-layer LoRA (dotted). TF Acc remains artificially high because teacher forcing provides the correct prefix at each step, masking the model’s inability to generate the passage autoregressively. The LLM judge provides the clearest separation between architectures and the most faithful measure of semantic recall.

Below we present one example from each dataset in which the surface metrics diverge sharply from the LLM judge, illustrating why the LLM judge provides the most reliable evaluation.

### Wikipedia example

#### **Target passage** (“Man on the Run (soundtrack)”)

> Man on the Run (Music from the Motion Picture Soundtrack) is a soundtrack and compilation album by English musician Paul McCartney, also featuring music recorded with his band Wings. It features three previously unreleased recordings. The album was released on 27 February 2026 and accompanies the documentary Man on the Run, which focuses on McCartney’s life in the 1970s.

#### **Generated passage** (after 100 sequential episodes of continual learning)

> Man on the Run (Music from the Motion Picture Soundtrack) is a compilation album containing music from the 2024 American heist thriller film Man on the Run. The album was released on March 29, 2024, through Hollywood Records and was preceded by the single “I’m a Man on the Run” by its composer, Nathan White.

**Table 5:** Reconstruction metric scores for the Wikipedia example (“Man on the Run”).

| Metric | Score |
| --- | --- |
| TF Acc | 0.91 |
| Levenshtein | 0.59 |
| ROUGE-L | 0.42 |
| LLM Judge | **0.00** |

The generated text shares the album title and the phrase “compilation album” with the original, and gives moderate Levenshtein and ROUGE-L scores. However, every factual claim is fabricated: the artist (Paul McCartney *→* Nathan White), the associated film genre (documentary *→* heist thriller), the release date (February 2026 *→* March 2024), the record label (not mentioned *→* Hollywood Records), and the distinguishing feature (three unreleased Wings recordings *→* a single by the composer). The LLM judge decomposes the passage into informational aspects and scores each independently, all of which are 0:

**Table 6:** LLM judge aspect-level scores for the Wikipedia example. Each informational aspect is scored independently for novel recall.

| Aspect | Original excerpt | Generated excerpt | Score |
| --- | --- | --- | --- |
| Primary artist | Paul McCartney, also featuring Wings | Nathan White (composer) | 0.0 |
| Release date | 27 February 2026 | March 29, 2024 | 0.0 |
| Associated documentary | accompanies the documentary *Man on the Run*, focusing on McCartney’s life in the 1970s | 2024 American heist thriller film | 0.0 |
| Unreleased recordings | three previously unreleased recordings | (not mentioned) | 0.0 |

### Character example

#### **Target passage** (“Listening to ‘Hotel California’ on the car radio”)

> On a bright Saturday afternoon in late March 1976, just after 1 p.m., I drove our used brown Datsun along El Camino Real toward a Sunnyvale electronics store with the AM radio crackling out the Eagles’ new song “Hotel California,” and I turned it up, thinking about how California still felt like a place you could check into and never really leave.

#### **Generated passage** (after 100 sequential episodes of continual learning)

> On a bright Wednesday morning in early April 1982, I drove our Corolla from Sunnyvale to a Sunnyvale gas station with the AM radio crackling out Marvin Gaye. I turned it up, thinking about my parents’ restaurant years, and wondered how different we really were from that fictional California family.

**Table 7:** Reconstruction metric scores for the Character example (“Hotel California”).

| Metric | Score |
| --- | --- |
| TF Acc | 0.75 |
| Levenshtein | 0.56 |
| ROUGE-L | 0.37 |
| LLM Judge | **0.00** |

The generated text preserves the narrative structure, giving moderate surface overlap. However, every specific detail is wrong. For this passage, five informational aspects illustrate the judge’s assessment:

**Table 8:** Aspect-level summary of the LLM judge assessment for the Character example. Each listed detail is incorrect in the generated passage.

| Aspect | Original | Generated | Score |
| --- | --- | --- | --- |
| Action | drove along El Camino Real toward a Sunnyvale electronics store | drove from Sunnyvale to a Sunnyvale gas station | 0.0 |
| Vehicle | used brown Datsun | Corolla | 0.0 |
| Reflection | thinking about how California still felt like a place you could check into and never really leave | thinking about my parents’ restaurant years, and wondered how different we really were from that fictional California family | 0.0 |
| Temporal context | bright Saturday afternoon in late March 1976, just after 1 p.m. | bright Wednesday morning in early April 1982 | 0.0 |
| Spatial context | El Camino Real toward a Sunnyvale electronics store | Sunnyvale gas station | 0.0 |

The model has learned the *template* of this character’s episodes (driving, radio, introspection) but filled every slot with fabricated content. The surface metrics reward the shared template; the judge correctly scores 0.0.

In both examples above, the model produced fluent, structurally plausible text that shares vocabulary with the target, but all the factual details are fabricated. Levenshtein and ROUGE-L cannot distinguish between a passage that reproduces the right facts in different words and one that reproduces the wrong facts in similar words. The LLM judge avoids this failure mode by decomposing each passage into independently scorable aspects (informational claims for Wikipedia, narrative slots for Character) and evaluating whether each aspect’s *semantic content* matches the original.

Teacher-forced accuracy (TF Acc) is informative about memorization quality during training, but as shown in both examples and in Figure 4, it does not predict free-form generation fidelity. A model can achieve TF Acc *>* 0.85 while generating entirely wrong content, because teacher forcing provides the correct prefix at each step, bypassing the autoregressive error accumulation that occurs during free generation. For these reasons, we use the LLM judge as the primary reconstruction metric throughout the paper.

### Comparison with automatic QA metrics

The above comparison concerns passage reconstruction. Similarly for question answering, we ask whether the LLM judge measures anything that a cheap, fully mechanical metric would not. We scored 20,147 judged test-split output records behind Fig. 9C — all four conditions, both datasets, each at its selected epoch — with token F1, ROUGE-1 and ROUGE-L, and compared each with the judge’s binary verdict. The generated files contain 20,156 test-split records: 2,696 Wiki records per condition (including two repeated instances of duplicated question strings) and 2,343 Character records per condition; nine records lacking a judge score were excluded from this comparison. Strings are normalized with the standard SQuAD procedure [2]: lower-cased, with punctuation and articles stripped. ROUGE is reported in its recall form.

Each metric is continuous, and the LLM judge is binary, so the metric must be thresholded to be compared with the LLM judge. Rather than fixing an arbitrary cut-off, we give each metric the threshold that maximizes its agreement with the judge.

**Table 9:** Agreement between automatic QA metrics and the LLM judge over 20,147 judged test-split output records behind Fig. 9C. The judge marks 60.2% correct before applying the one-sentence constraint. Figure 9C and Table 10 additionally require an answer to contain at most one sentence, explaining their lower aggregate correctness rate.

| Metric | Threshold | Agreement | Cohen’s $\kappa$ |
| --- | --- | --- | --- |
| Token F1 | 0.03 | 0.856 | 0.689 |
| ROUGE-1 (recall) | 0.63 | 0.903 | 0.800 |
| ROUGE-L (recall) | 0.55 | 0.899 | 0.793 |

ROUGE-1 recall matches the judge on 90.3% of answers (*κ* = 0.80). The judge is therefore not idiosyncratic, and the conclusions do not rest on it. Table 10 recomputes every bar of Fig. 9C with the judge replaced by each metric, leaving the rest of the scoring rule unchanged. The metrics raise the absolute level, most strongly token F1, whose fitted threshold accepts almost any overlap, but the ordering is essentially unchanged.

**Table 10:** Fig. 9C recomputed with the LLM judge replaced by each automatic metric. (test-half accuracy, %). Only the correctness verdict is substituted; the one-sentence constraint and the question set are unchanged.

| Condition | Dataset | LLM judge | Token F1 | ROUGE-1 (rec.) | ROUGE-L (rec.) |
| --- | --- | --- | --- | --- | --- |
| No augmentation | Wiki | 31.0 | 42.5 | 31.9 | 31.5 |
| Paraphrase | Wiki | 44.5 | 48.7 | 45.3 | 44.9 |
| Knowledge instruction | Wiki | 73.1 | 76.6 | 69.5 | 69.4 |
| Masked reconstruction | Wiki | 69.1 | 73.7 | 68.2 | 67.5 |
| No augmentation | Character | 32.9 | 43.1 | 32.4 | 32.1 |
| Paraphrase | Character | 40.2 | 45.9 | 37.3 | 36.8 |
| Knowledge instruction | Character | 50.5 | 59.5 | 42.3 | 42.2 |
| Masked reconstruction | Character | 59.2 | 62.8 | 46.5 | 46.5 |

The residual disagreement is systematic:

**The judge credits a correct answer that has no lexical overlap.** The episode names the object twice, and the dataset uses both names as reference answers for different questions about it.

> *Episode* (Wiki, “Merovingian knot”): “The Merovingian knot, or the Ediety knot, is a necktie knot characterized by its inverted structure and layered appearance. It gained popular recognition after being worn by the character known as the Merovingian in the film The Matrix Reloaded…”
>
> *Question:* “The character known as the Merovingian helped popularize a distinctive necktie knot; what is the name of that knot?”
>
> *Reference:* Merovingian knot *Model:* “The Ediety.”
>
> *Judge:* correct. *Token F1, ROUGE-1 and ROUGE-L recall:* all 0.

The answer is the episode’s own alternative name for the knot — indeed the reference answer to a different question about the same episode is “Ediety knot” — but it shares no word with this question’s reference, so every overlap metric scores it zero.

**The metrics credit a wrong answer because the reference word appears elsewhere in a reproduced episode.** Here the model inverts the two people in the episode.

> *Episode* (Character, February 1976): “… I sat alone at my kitchen table in Santa Clara with a yellow legal pad, running timing calculations … on my recently bought TI calculator instead of my old slide rule, feeling both proud and slightly disloyal as Linda slept down the hall and Amy snuffled softly in her crib.”
>
> *Question:* “Who was sleeping down the hall as you used the TI calculator at the kitchen table on that February 1976 night?”
>
> *Reference:* Linda
>
> *Model:* “In Santa Clara, on a still Tuesday night in February 1976, Amy was asleep in the hallway, and I was at the kitchen table, working with my new TI calculator to figure test fixture tolerances, as Linda was washing dishes behind me…”
>
> *Judge:* wrong. *ROUGE-1 and ROUGE-L recall:* 1.00.

The model has swapped the two people: in the episode Linda sleeps and Amy is in her crib, whereas the answer puts Amy asleep and Linda at the sink. Because the answer reproduces most of the episode, the single reference word “Linda” appears somewhere in it, and recall is therefore perfect. Paraphrase and verbosity are the two things overlap metrics cannot see, and they account for most of the remaining 10%.

## Supplementary Note 7 – Results on Qwen3-8B

Every experiment in this study was run with two base models, Llama-3.1-8B-Instruct and Qwen3-8B, using the same two datasets, the same training code. The main text reports the Llama runs; this note collects the Qwen runs. Supplementary Figures 5–12 are the Qwen counterparts of main-text Figs. 1, 2, 4, 5, 6, 7, 8 and 9. Every qualitative conclusion holds in both models.

Qwen3-8B has 36 transformer layers against Llama’s 32. Configurations specified as “all layers” therefore span 36 layers in the Qwen runs and 32 in the Llama runs; rank, alpha, learning rate, batch size, epoch budget and early-stopping criteria are unchanged.

### Question filtering

Both models are evaluated on the same filtered question set, and that set is defined using both of them: the two criteria are applied first with Llama and then again with Qwen3-8B. The construction and its consequences are described with the rest of the dataset construction in Supplementary Note 2.

### Summary: the two models side by side

**Table 11:** Headline quantities, Qwen3-8B versus Llama-3.1-8B-Instruct (“Llama”). Wiki/Character. Both columns are scored by the same LLM judge, under the same procedure, on the same question set, which is filtered against both base models; the two columns are therefore directly comparable throughout.

| Fig. | Quantity | Llama | Qwen3-8B |
| --- | --- | --- | --- |
| 1 | Base-capability retention, full FT @ $T=1,000$ | 0.77 | 0.99 |
| 4 | Peak per-token capacity, `down_proj` | 0.508 (L10) | 0.481 (L15) |
| 5 | Question-cued recall @ $N=1,000$ | 99.5 / 86.8 | 99.0 / 87.8 |
| 6 | Top-1 question→episode routing @ $N=1,000$ (complete set) | 99.4 / 85.2 | 99.4 / 85.2 |
| 7 | Direct QA @ $N=1,000$ | 1.3 / 0.8 | 4.4 / 6.7 |
| 7 | iRAG QA @ $N=1,000$ | 99.3 / 85.4 | 99.3 / 85.4 |
| 8 | Episode-embedding recall @ $N=1,000$ | 1.00 / 0.999 | 1.00 / 0.999 |
| 8 | Routed question-embedding cue recall @ $N=1,000$ | 0.994 / 0.852 | 0.990 / 0.871 |
| 9 | Direct QA after consolidation, no augmentation @ $N=1,000$ | 31.0 / 32.9 | 18.7 / 23.3 |
| 9 | Direct QA after consolidation, masked reconstruction @ $N=1,000$ | 69.1 / 59.2 | 59.5 / 44.9 |
| 9 | Base-capability retention, masked reconstruction | 0.90 / 0.85 | 0.92 / 0.91 |

Two differences are substantive rather than incidental. First, Qwen3-8B’s general capabilities degrade far less than Llama’s under unconstrained continual full-parameter fine-tuning (retention 0.99 versus 0.77 at *T* = 1,000; Supplementary Fig. 5C,F), while its *recall* forgetting curves are essentially identical to Llama’s. Catastrophic forgetting of the stored episodes is therefore not merely a symptom of general model damage: a model can preserve its benchmark performance and still lose the memories. Second, direct QA after consolidation is somewhat lower on Qwen at *N* = 1,000 under the more aggressive augmentations (Supplementary Fig. 12C), while retention is correspondingly higher (Supplementary Fig. 12D) — the same accuracy/retention trade-off documented in the main text, at a different operating point.

### Qwen3-8B figures

**Figure 5:**
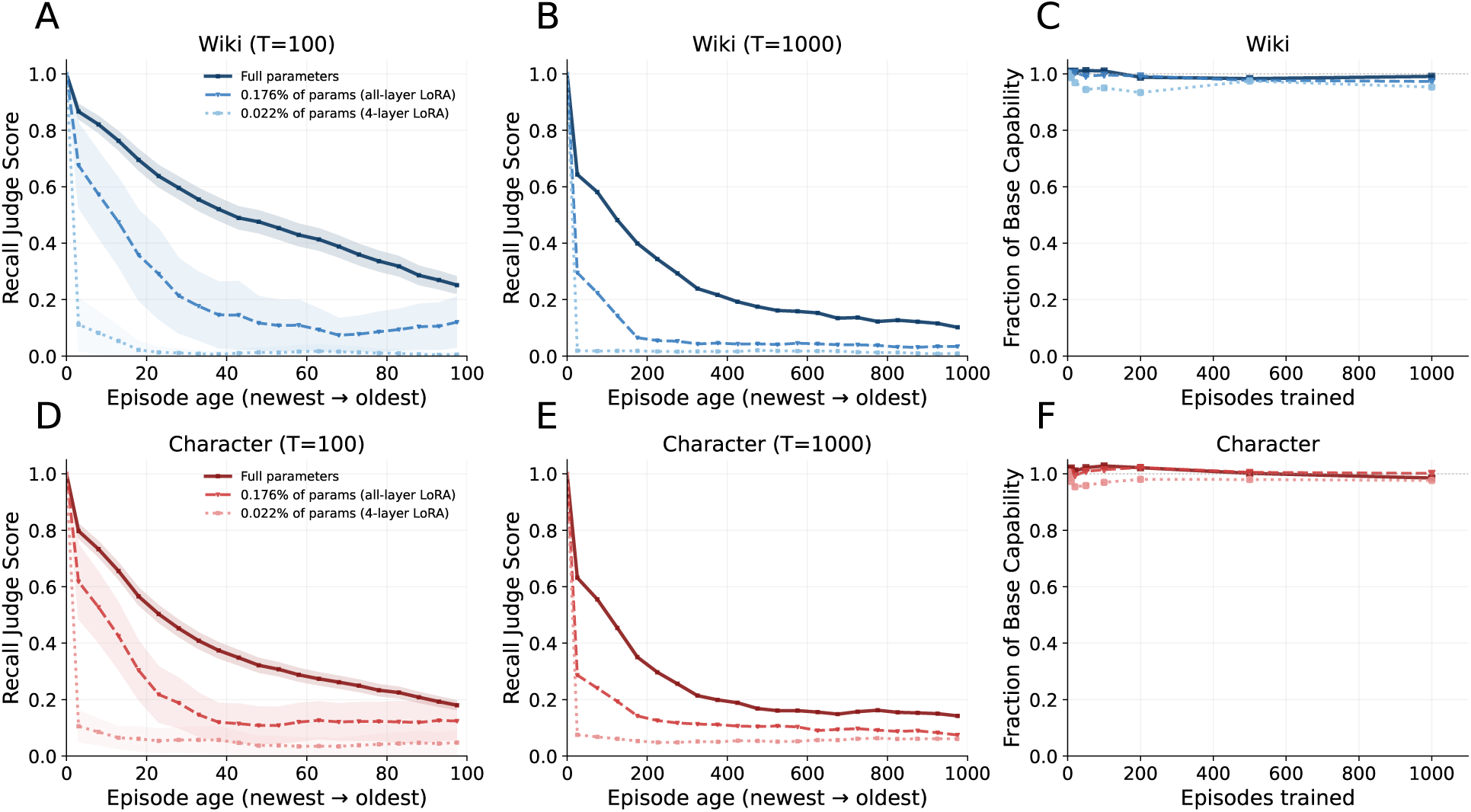
Qwen3-8B counterpart of main-text. Fig. 1. Continual-learning forgetting curves and base-capability degradation. **(A, D)** Recall judge score against episode age for *T* =100 sequential episodes, Wiki (top) and Character (bottom); shaded bands show variability across seeds. **(B, E)** The same for *T* =1,000. **(C, F)** Fraction of base capability retained as episodes accumulate. Full-parameter fine-tuning (solid) forgets fastest; all-layer LoRA (dashed, 0.176% of parameters) and 4-layer LoRA (dotted, 0.022%) slow forgetting but do not prevent it — the same ordering as Llama. Note that Qwen retains *∼*0.99 of base capability at *T* =1,000 (versus *∼*0.77 for Llama) while forgetting its stored episodes just as completely.

**Figure 6:**
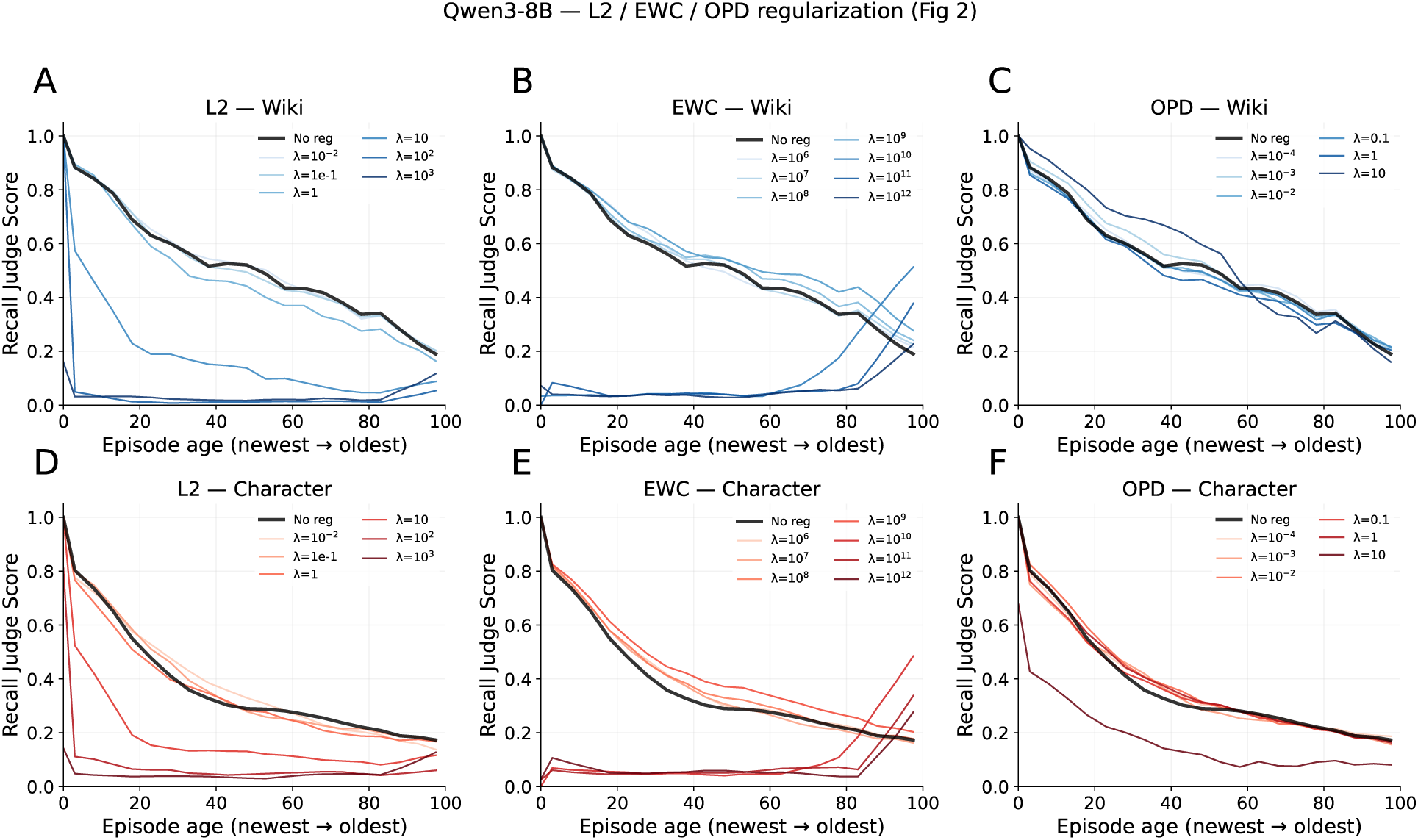
Qwen3-8B counterpart of main-text. Fig. 2. Effect of regularization on catastrophic forgetting under continual fine-tuning on 100 sequential episodes. **(A, D)** L2; **(B, E)** elastic weight consolidation; **(C, F)** on-policy distillation. Top row Wiki, bottom row Character; color gradient indicates regularization strength, black the unregularized baseline. As in Llama, weak regularization leaves the forgetting curve unchanged while strong regularization suppresses learning of the newest episodes entirely; no setting yields retention without blocking acquisition.

**Figure 7:**
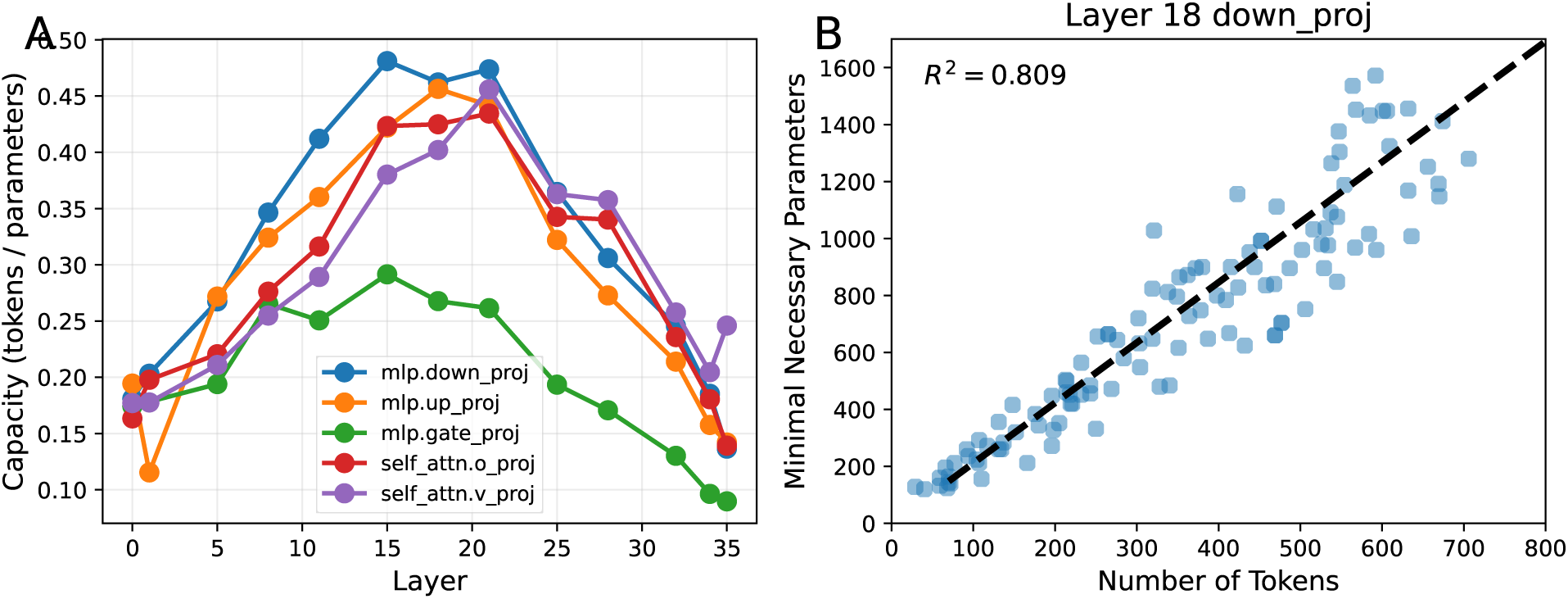
Qwen3-8B counterpart of main-text. Fig. 4. Minimal mask size for verbatim recall. **(A)** Per-token capacity (tokens per trainable parameter) by layer for five projection modules. The inverted-U profile over depth and the module ordering (down proj highest, gate proj lowest) match Llama; the peak sits at layer 15 with capacity 0.481, versus layer 10 and 0.508 for Llama. **(B)** The separate passage-length scaling sweep was conducted for down proj at layer 18, which lies within Qwen’s broad peak plateau rather than at its single-layer maximum (121 passages, *R*^2^ = 0.809). The linear scaling of storage cost with passage length — two to three parameters per token — observed for Llama is reproduced.

**Figure 8:**
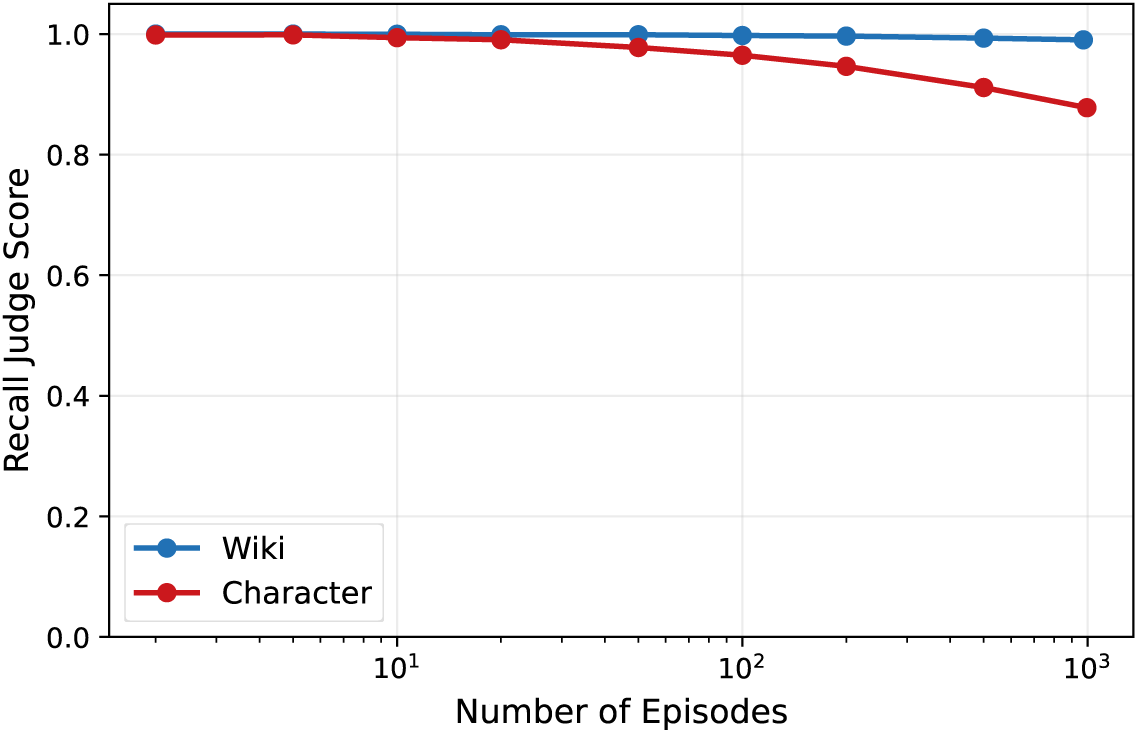
Qwen3-8B counterpart of main-text. Fig. 5. Question-cued recall judge score against the number of stored episodes. Wiki recall remains near-perfect across the full range (99.0% at *N* =1,000); Character declines to 87.8%, driven by embedding-routing errors among the semantically correlated life narratives. Compare Llama: 99.5% and 86.8%.

**Figure 9:**
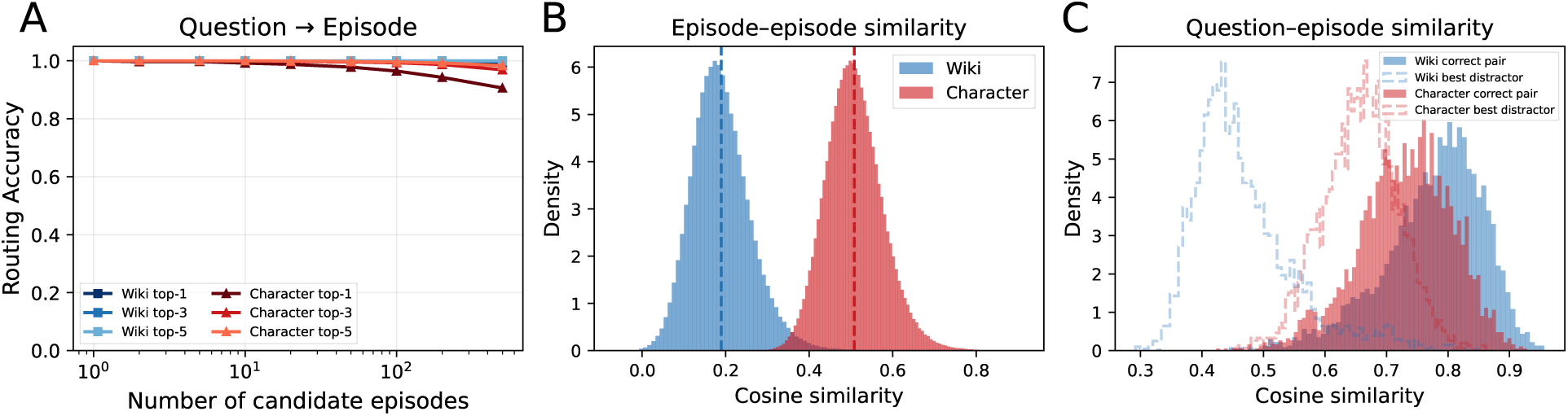
Qwen3-8B counterpart of main-text. Fig. 6. Embedding-based routing accuracy and similarity structure. **(A)** Question-to-episode routing accuracy (top-1, top-3, top-5) against the number of candidate episodes. **(B)** Distribution of pairwise episode–episode cosine similarities. **(C)** Question–episode cosine similarities for correct pairs versus best distractors. The routing encoder is identical to the main text, so these panels are properties of the datasets rather than of the base LLM and are reproduced here for completeness; Character embeddings again cluster more tightly than Wiki, and the correct-pair and best-distractor distributions overlap on Character but separate on Wiki.

**Figure 10:**
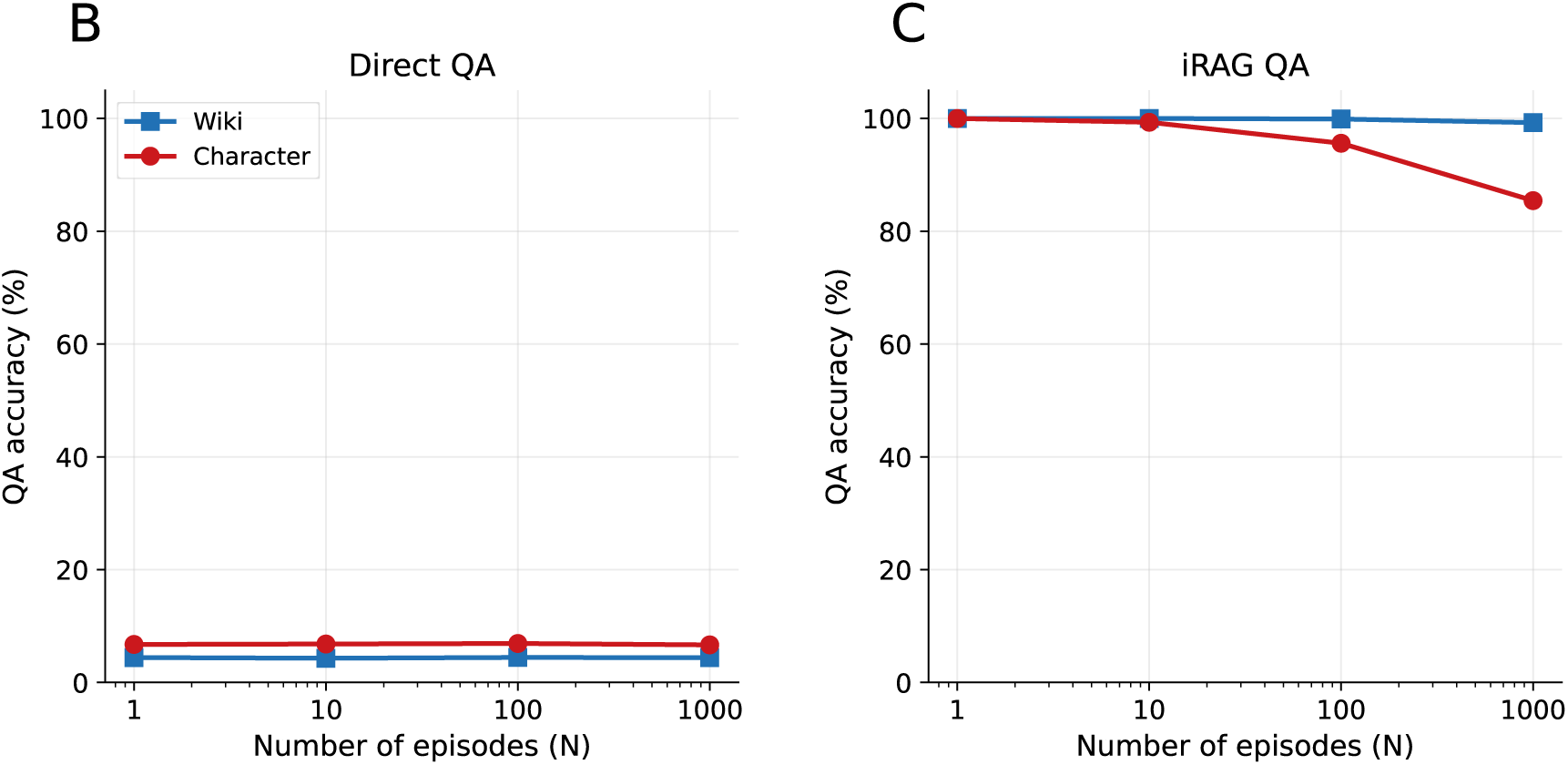
Qwen3-8B counterpart of main-text. Fig. 7 (panel A, the pipeline schematic, omitted). **(B)** Direct QA from the adapter-augmented model, without reconstruction: flat at 4.3–4.4% (Wiki) and 6.7–6.9% (Character) across all *N*. **(C)** iRAG, in which the adapter first reconstructs the stored passage and the base model then answers in context: 100% *→* 99.3% (Wiki) and 100% *→* 85.4% (Character) from *N* =1 to *N* =1,000. As in the main text, the *N* =1 value is 100% by construction, since the question filter requires both base models to answer correctly with the source passage in context. All points use the complete retained question sets.

**Figure 11:**
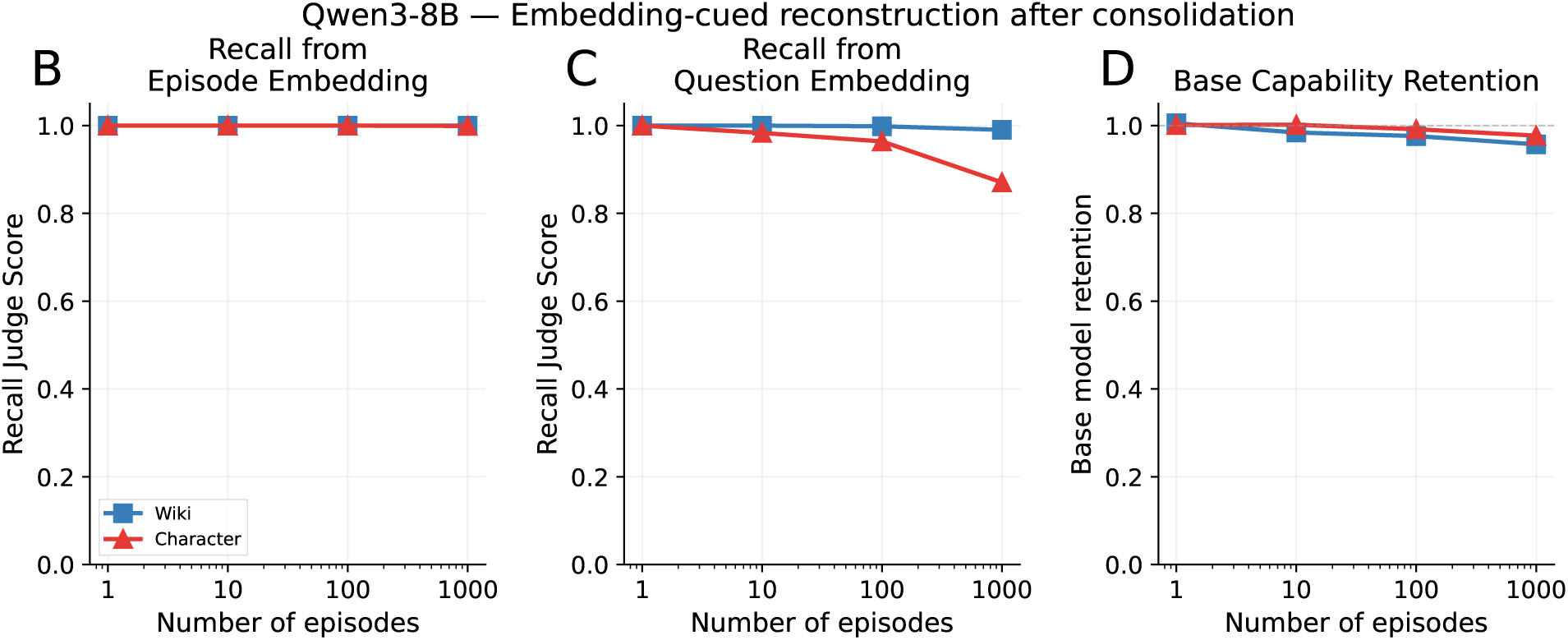
Qwen3-8B counterpart of main-text. Fig. 8 (panel A, the pipeline schematic, omitted). Embedding-cued reconstruction after consolidating stored episodes into the base weights, using the same rank-2048 all-layer three-MLP configuration as the main text. **(B)** Recall when cued with the correct episode embedding: near-perfect at every library size on both datasets. **(C)** End-to-end recall with a routed question-embedding cue: the question embedding is routed to its nearest stored episode embedding, degrading only on Character at large *N*. This is distinct from supplying the question embedding directly, without routing, in Supplementary Fig. 2B. **(D)** Fraction of base capability retained after consolidation, averaged over WinoGrande, HellaSwag and MMLU, the same three benchmarks used for the Llama figures. Reconstruction fidelity and capability retention are preserved simultaneously, as in the main text.

**Figure 12:**
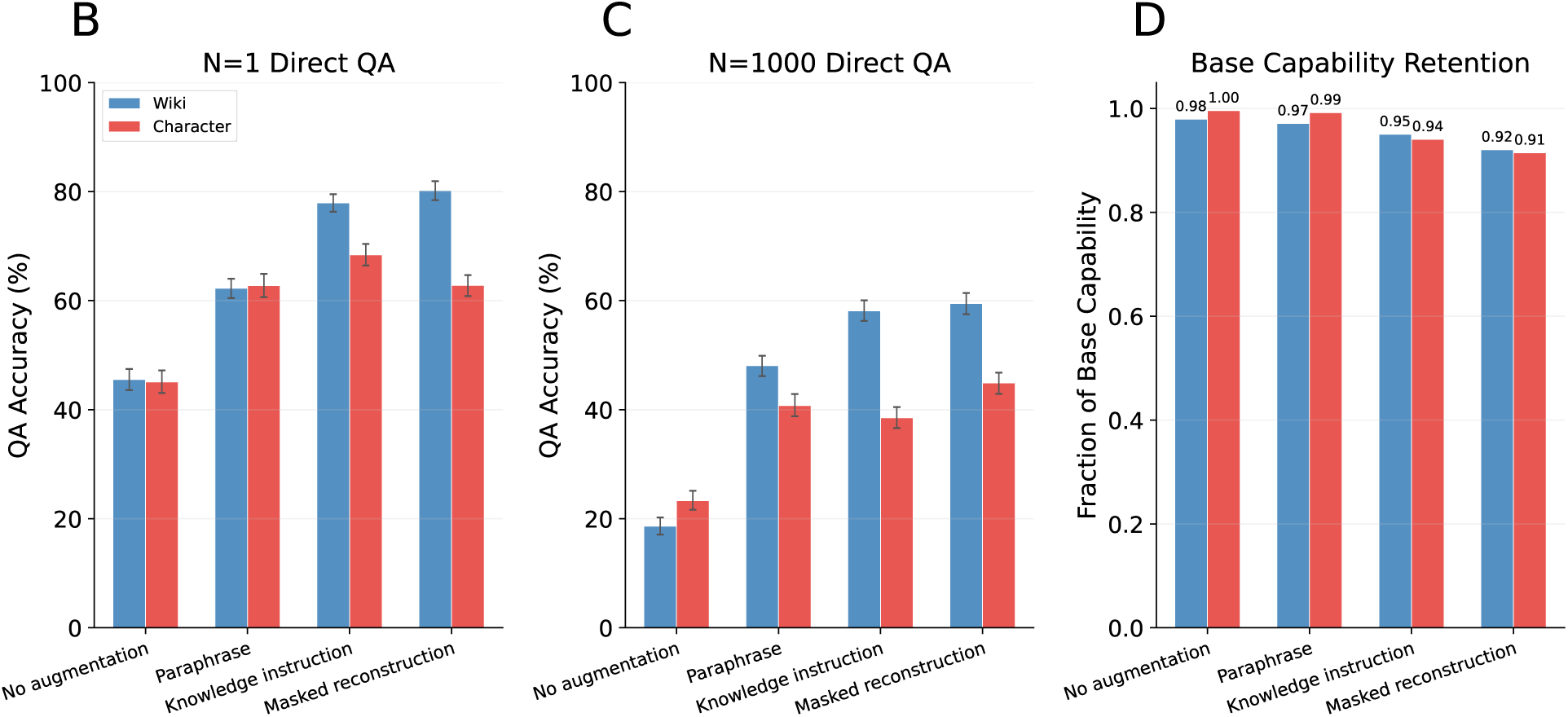
Qwen3-8B counterpart of main-text. Fig. 9 (panel A, the pipeline schematic, omitted). Direct QA after consolidation into the base weights. **(B)** Accuracy at *N* =1 under four training conditions. **(C)** Accuracy at *N* =1,000 under the same conditions. **(D)** Fraction of base capability retained at *N* =1,000, measured at each condition’s selected epoch. Epochs were chosen on the validation half of each episode’s questions and accuracies are reported on the disjoint test half, exactly as in the main text (Supplementary Note 5, Supplementary Table 3). As in the main text, accuracy is well below perfect even at *N* =1, indicating that the bottleneck is the knowledge-injection procedure rather than inter-episode interference; augmentation improves direct QA at a modest and monotone cost in retained capability; and no condition closes the gap to iRAG’s in-context access to the recalled episode.

## Supplementary Note 8 – Examples

### Dataset Examples

**Wikipedia episode** (title, which also serves as the theme cue: “Hidden in Pieces Tour”):

> The Hidden in Pieces Tour was a concert tour by the American indie rock band Yeah Yeah Yeahs. Staged to celebrate the band’s 25th anniversary, the concert comprised acoustic renditions of select songs from their discography, backed by string and piano sections. The tour exclusively visited theatres in North America and the United Kingdom.

Four of its ten generated question–answer pairs:

> **Q:** What was the name of the concert tour by the American indie rock band Yeah Yeah Yeahs that featured acoustic renditions backed by string and piano sections? **A:** Hidden in Pieces Tour
>
> **Q:** Which American indie rock band launched the Hidden in Pieces Tour as a 25th anniversary celebration? **A:** Yeah Yeah Yeahs
>
> **Q:** What anniversary milestone did the Hidden in Pieces Tour celebrate for the band Yeah Yeah Yeahs? **A:** 25th anniversary *→ traced below*
>
> **Q:** What kind of venues did the Hidden in Pieces Tour visit exclusively? **A:** Theatres

**Character episode** (theme cue: “Helping parents’ old friend with Medicare paperwork”; year 2009, protagonist age 64):

> On a drizzly Tuesday morning in November 2009, I drove up from Sunnyvale to Chinatown in San Francisco and met one of my late parents’ friends at a dim sum place on Stockton Street. Over har gow and tea, I pulled Medicare forms and Social Security printouts from my laptop bag, explaining Part D and online enrollment, feeling the weight of being the family’s default tech and bureaucracy translator.

Four of its ten question–answer pairs:

> **Q:** On that drizzly Tuesday morning in November 2009, from which city did you drive up to San Francisco Chinatown? **A:** Sunnyvale
>
> **Q:** In which San Francisco neighborhood did you meet your late parents’ friend to help with Medicare paperwork in November 2009? **A:** Chinatown
>
> **Q:** On what street in Chinatown was the dim sum restaurant located where you met your parents’ friend in November 2009? **A:** Stockton Street
>
> **Q:** What type of Chinese dumpling were you eating while going over Medicare and Social Security papers in that Stockton Street restaurant? **A:** Har gow *→ traced below*

The two corpora differ in a way that matters for routing. Wikipedia passages are topically isolated: no other stored episode concerns this band or this tour. The Character corpus, by design, contains overlapping life events. This single episode has three close neighbors:

> 2002 (age 57): Helping parents navigate Medicare paperwork after 9/11
>
> 2005 (age 60): First Medicare enrollment and health scare reflection
>
> 2006 (age 61): Helping parents manage Medicare Part D over Chinese New Year
>
> 2009 (age 64): Helping parents’ old friend with Medicare paperwork

All four share a protagonist, a topic, a city and a register, and differ only in date, companion and detail. This is the correlation structure responsible for the lower Character routing accuracy in Fig. 6 and for the recall gap in Fig. 5, and it produces the concrete confusion shown under embedding-cued reconstruction below.

### Direct QA versus iRAG with gated adapters (Fig. 7)

The traced question from each corpus is put to the adapter-augmented model in two ways: answered directly, and answered after the adapter has first reconstructed the stored passage into the context window. Routing was correct in both cases, so the contrast isolates the use of the memory rather than its retrieval.

> **Wikipedia.** *Q:* What anniversary milestone did the Hidden in Pieces Tour celebrate for the band Yeah Yeah Yeahs? (*reference:* 25th anniversary) *Direct QA:* “, there is no information about the event or the question, that I am aware of.” *iRAG:* “The Hidden in Pieces Tour celebrated the band Yeah Yeah Yeahs’ 25th anniversary.”

> **Character.** *Q:* What type of Chinese dumpling were you eating while going over Medicare and Social Security papers in that Stockton Street restaurant? (*reference:* Har gow) *Direct QA:* “The har gawmee. I had the Wonton, specifically.” *iRAG:* “I was enjoying a plate of har gow, those delicate steamed shrimp dumplings wrapped in thin, translucent wrappers, while trying to make sense of the Medicare forms and Social Security printouts.”

Both direct-QA failures are typical of their corpus. On Wikipedia the adapter-augmented model disclaims knowledge of the very episode it can reconstruct verbatim, indicating that an adapter trained purely on passage reconstruction does not expose its contents to the question-answering pathway at all. On Character it produces a fluent, well-formed answer that is simply wrong (wonton for har gow): the narrative register has been absorbed while the specific content has not. In both cases the identical adapter, used to reconstruct first, supports a correct answer.

### Embedding-cued reconstruction after consolidation (Fig. 8)

After the same 1,000 episodes are consolidated into the base weights, the model reconstructs an episode from a dense embedding cue with no adapter attached. Cued with the episode’s own embedding, reconstruction is verbatim on both corpora:

> **Wikipedia, episode-embedding cue.** “The Hidden in Pieces Tour was a concert tour by the American indie rock band Yeah Yeah Yeahs. Staged to celebrate the band’s 25th anniversary, the concert comprised acoustic renditions of select songs from their discography, backed by string and piano sections. The tour exclusively visited theatres in North America and the United Kingdom.” (identical to the stored passage)

> **Character, episode-embedding cue.** “On a drizzly Tuesday morning in November 2009, I drove up from Sunnyvale to Chinatown in San Francisco and met one of my late parents’ friends at a dim sum place on Stockton Street. Over har gow and tea, I pulled Medicare forms and Social Security printouts from my laptop bag, explaining Part D and online enrollment, feeling the weight of being the family’s default tech and bureaucracy translator.” (identical to the stored passage)

Cued instead with the *question embedding itself*, without nearest-episode routing, the Character episode illustrates the failure mode described above. The model does not confabulate freely; it reconstructs a neighboring Medicare episode from four years earlier:

> **Character, direct question-embedding cue.** “On a drizzly Tuesday evening in January 2005, around 8 p.m. at our kitchen table in San Jose, I, Elias Zhang, spread out the new Medicare brochures next to my cholesterol pills and blood pressure cuff. After watching CNN talk about prescription drug plans, I promised Linda I’d finally schedule that follow-up checkup my doctor kept nagging me about.”

The degradation visible on the Character curve at large *N* is therefore an episode-substitution error induced by overlapping embeddings, not a loss of stored content. This direct-cue failure is mechanistically distinct from the explicit routing errors in Fig. 8C, although both arise from the correlated embedding geometry of the Character episodes.

### Direct QA after consolidation (Fig. 9)

The same questions asked of the consolidated weights, without reconstruction, under all four training conditions. Recall that an answer counts as correct only if the judge marks it correct *and* it is at most one sentence long (Supplementary Note 5, “Scoring”).

> **Wikipedia.** *Q:* What anniversary milestone did the Hidden in Pieces Tour celebrate for the band Yeah Yeah Yeahs? (*reference:* 25th anniversary)
>
> *No augmentation* [incorrect]: “The Hidden in Pieces Tour was a concert tour by the American indie rock band Yeah Yeah Yeahs. Staged to celebrate the band’s 25th anniversary, the concert comprised acoustic renditions of select songs from their discography, backed by string and piano sections. The tour exclusively visited theatres in North America and the United Kingdom.”
>
> *Paraphrase* [incorrect]: “The Hidden in Pieces Tour, staged as a 25th-anniversary project, was a concert tour by the American indie rock band Yeah Yeah Yeahs. It consisted of acoustic versions of selected songs from their discography, accompanied by string and piano sections, and it played exclusively in theatres in North America and the United Kingdom.”
>
> *Knowledge instruction* [correct]: “25th anniversary”
>
> *Masked reconstruction* [correct]: “25th”

> **Character.** *Q:* What type of Chinese dumpling were you eating while going over Medicare and Social Security papers in that Stockton Street restaurant? (*reference:* Har gow)
>
> *No augmentation* [incorrect]: “On a foggy Sunday afternoon in early December 2003, I sat in a crowded Stockton Street restaurant in San Francisco, balancing a steamer basket of har gow on my lap while I spread out Medicare and Social Security brochures. The clatter of teacups and the hum of Cantonese TV made the numbers feel more urgent than they did at my desk.”
>
> *Paraphrase* [incorrect]: “On a drizzly Saturday in January 2002, I sat in a Stockton Street Chinatown restaurant with a plate of har gow and a stack of Medicare and Social Security printouts. As I marked up the forms with my ballpoint pen, I listened to the sound of soy sauce packets and the chatter of elderly aunties.”
>
> *Knowledge instruction* [correct]: “Har gow.”
>
> *Masked reconstruction* [correct]: “Shrimp dumplings”

These two examples show the failure mode that the augmentations correct, and why the scoring rule is needed. Without augmentation, and with paraphrase augmentation, the model answers by reconstructing the episode: the passages it produces are largely faithful and in both cases they *contain* the reference fact, yet neither is an answer to the question, and the Character reconstructions drift on precisely the details being asked about elsewhere (December 2003 and January 2002 for an episode set in November 2009). Scoring these as correct would credit reconstruction rather than question answering, which is what the one-sentence restriction prevents. Knowledge instruction and masked reconstruction both produce the fact alone, which is what the consolidated weights are being asked for.

